# Palbociclib CDK4/6- and Crizotinib MET/ALK/ROS1-inhibitors Synergize to Enhance Senescence and Immune Recognition in Melanoma Cells Independently of *BRAF*/*NRAS* Status

**DOI:** 10.1101/2025.07.04.663080

**Authors:** Fan Zhang, Lola Boutin, Ishani Das, Jeroen Melief, Madhurendra Singh, Marina Stantic, Lu Zhang, Mohammad Alzrigat, Alireza Azimi, Lucas Baldran, Wesam Bazzar, Michelle Da Silva Liberio, Jacob Goodwin, Rainer Tuominen, Veronica Höiom, Fredrik Jerhammar, Suzanne Egyhazi Brage, Johan Hansson, Rolf Kiessling, Galina Selivanova, Klas G. Wiman, Margareta Wilhelm, Lars-Gunnar Larsson

## Abstract

“Pro-senescence therapy”, which triggers both permanent cell cycle arrest and an immune response, has been proposed as a strategy for cancer treatment, but is still controversial. To assess this strategy in melanoma, we performed a high throughput microscopy-based senescence screen utilizing a panel of melanoma cell lines with different driver mutations and a collection of clinical and experimental drugs. We found that vemurafenib and trametinib, which inhibit BRAF^V600E^ and MEK1/2, respectively, induced senescence in some but not all BRAF-mutant cell lines. In contrast, palbociclib, BKM-120 and crizotinib, which inhibit CDK4/6, PI3K, and MET/ALK/ROS1, respectively, triggered senescence in most cell lines, irrespective of BRAF/NRAS mutation status, and overcame intrinsic and acquired vemurafenib resistance. The combination of palbociclib and crizotinib synergized to further enhance the senescence response in all cell lines irrespective of BRAF/NRAS mutation status, increased the expression of SASP factors, such as IL-1α and β, and HLA class I and other markers for recognition by NK and T cells. Further, this combination caused a significant increase in CD8+ T cells and pro-inflammatory macrophages in the tumor microenvironment and a marked reduction of mouse melanoma tumor growth that was dependent on CD8+ T cells, suggesting increased immune surveillance. Our findings suggest that pro-senescence therapy based on concomitant inhibition of both CDK4/6 and MET/ALK/ROS1 could be developed further as an alternative treatment strategy for melanoma.

Significance: Pro-senescence therapy based on combined targeting of CDK4/6 with palbociclib and MET/ALK/ROS1 with crizotinib inhibits melanoma tumor growth through anti-tumor immune response activation, providing an alternative treatment strategy for melanoma.

## Introduction

The incidence of cutaneous melanoma is increasing at one of the fastest rates among all cancer types, and it remains the skin cancer with highest mortality rate, causing around 55 000 deaths per year worldwide (1). Approximately 50% of advanced cutaneous melanomas carry a *BRAF* mutation, most commonly *BRAF^V600E^*, while about 25% harbor mutations in *NRAS* (2,3); both genes are upstream regulators of the MAP kinase pathway. The development of specific inhibitors targeting the mutated BRAF^V600E^ kinase and selective inhibitors of the downstream MEK1/2 kinases has drastically improved treatment outcome of advanced melanoma for more than a decade (4–6). A key feature of melanoma is its very high load of somatic mutations (2) resulting in neoantigens that can be recognized by the immune system. Consequently, the introduction of immune checkpoint inhibitors (ICI) blocking PD-1, CTLA-4 or other checkpoint molecules which stimulate T-cell engagement in the anti-tumor immune response was another major breakthrough. This has significantly improved survival in patients with advanced melanoma over the past 15 years (7,8). However, BRAF/MEK therapies are only effective in 50% of cases with the BRAF^V600E^-mutation, and beneficial effects with immunotherapies have also been limited to around 50% of the cases (1). In addition, most metastatic cancers eventually develop resistance to targeted therapies, leaving a large portion of advanced melanoma with limited treatment options (1,9), emphasizing the urgent need for new therapeutic strategies.

During the last decade, the concept of “therapy-induced senescence” (TIS) or “pro-senescence therapy” has emerged as an alternative strategy to combat cancer (10–12). Cellular senescence - defined as permanent cell cycle arrest - is along with apoptosis a main barrier against tumor development (13). It also plays an important role during embryogenesis and in adults during tissue repair and renewal (14). Senescence is triggered by different types of stress, e.g. aberrant oncogene expression (oncogene-induced senescence (OIS)), telomere dysfunction, oxidative and endoplasmic reticulum (ER) stress and DNA damage caused by for instance chemotherapeutic drugs or irradiation (10,12,15,16). As an example of OIS, oncogenic BRAF is a potent inducer of senescence in premalignant melanocytes in nevi of the skin, while progression to melanoma is accompanied by evasion of senescence (17,18). The senescence process is usually driven by two major tumor suppressor pathways, i.e. p53/p21 and p16/pRB pathways (10,12,19). However, there is a variety of senescence phenotypes described, depending on the senescence trigger, cell type, and microenvironment (20–22). An important aspect of the senescent phenotype is the senescence-associated secretory phenotype (SASP) (23,24). The biological function of SASP is to maintain the senescent state and to induce innate and adaptive immune responses leading to clearance of senescent cells, including senescent tumor cells (10,12,19,21,25,26). On the other hand, there is also multiple evidence suggesting that SASP components can promote tumor progression by stimulating inflammation, epithelial to mesenchymal transition (EMT), stemness, immune suppression, and other tumor-promoting processes, thereby undermining senescence-induced tumor suppression (10,12,21,25,27,28). In addition, tumor cells can escape senescence due to for instance loss/inactivation of *TP53*, *CDKN2A* (encoding p16 and p19ARF), *PTEN* or other tumor suppressors (3), or by activation of *MYC* (18,29,30).

Senescence and SASP can therefore be viewed as a double-edged sword in cancer development and progression. To overcome the negative sides of senescence, many efforts during recent years have attempted to introduce senolytics, i.e. substances that kill senescent cells, and senomorphics, i.e. substances that promote the “good” and prevent the “bad” sides of senescence, in order to promote immune surveillance of senescent cells and/or prevent senescence escape of tumor cells (10,12,31).

To explore the potential role of pro-senescence therapy in melanoma treatment, we performed a drug screen on a panel of melanoma cell lines representing the most common driver mutations and copy number alterations in human melanoma, including *BRAF^V600E^*, *NRAS^Q61R^*, *CDKN2A*, *PTEN*, *TP53*, *CDK4*, and *CCND1*. We show here that several of the drugs tested, either alone or in combination, particularly palbociclib combined with crizotinib, induce senescence and upregulate immune markers known to promote immune clearance in a broad range of melanoma cells. Further, using an immunocompetent melanoma mouse model we demonstrate that combined treatment with palbociclib and crizotinib stimulates infiltration of CD8+ T cells and M1-like macrophages and reduces tumor growth in a CD8+ T cell-dependent manner.

## Materials & Methods

### Cell culture

The human melanoma cell lines used are listed in Table 1. A375, SKMEL2, SKMEL28 and YUMM1.7 cells were purchased from the American Type Culture Collection (ATCC). The ESTDAB cell lines were obtained from European searchable Tumor cell Bank and database (ESTDAB), Germany (http://www.ebi.ac.uk/ipd/estdab). The early-passage KADA melanoma cell line was established from a stage III melanoma patient undergoing treatment in the oncology clinic at Karolinska University Hospital (32). A375-VR4 is a vemurafenib-resistant subline derived from A375 cells (33). Media for different melanoma cell lines, listed in Supplementary Table 1B, was supplemented with 10% fetal bovine serum (FBS) (Gibco), 2 mM L-glutamine, 50 U/ml of penicillin and 50 mg/ml of streptomycin (ThermoScientific). The YUMM1.7 mouse melanoma cell line (34) was cultured in DMEM: F12 GlutaMAX™ supplement (Thermo Fisher), supplemented with 10% FBS, 1% penicillin and streptomycin, and 1X MEM Non-Essential Amino Acids (Thermo Fisher). Cells were maintained in an incubator at 37 °C with 5% CO_2_ and optimal humidity. All cell lines used were tested negative for mycoplasma.

**Table 1.**
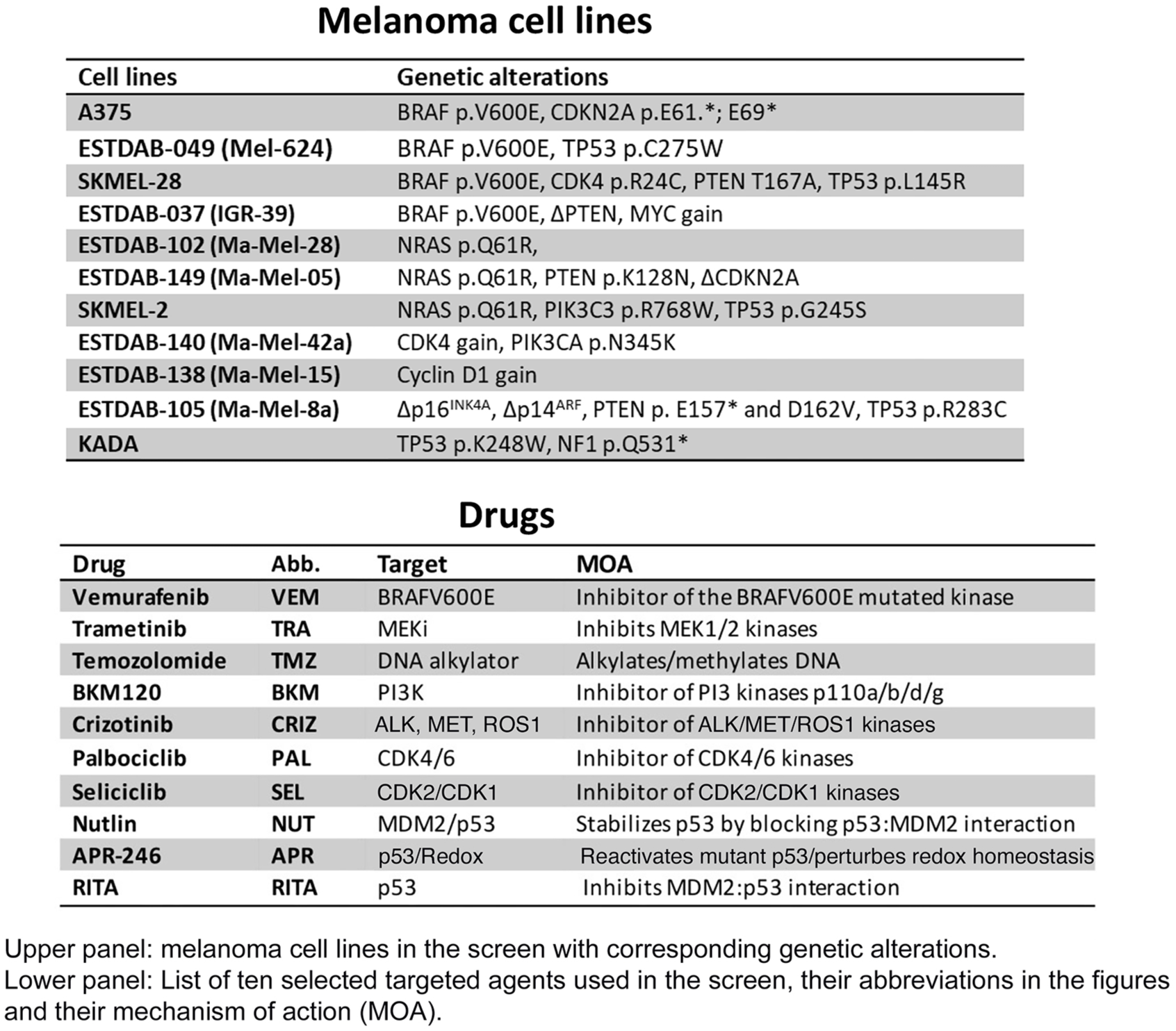
Melanoma cell lines and drugs used in the senescence screen Melanoma cell lines.

### Drugs and compounds

The drugs and compounds, the concentrations used, and their sources are listed in Table 1, Supplemental Table 1A and Supplemental Table 2.

### Whole genome sequencing

DNA was extracted from all cell lines using Allprep universal kit. This DNA was quantified using NanoDrop 2000 instrument and 100 ng was subjected to whole genome sequencing (WGS) using library build-up with the Nextera DNA library prep, Illumina platform and in-house developed post-read filtering (Science for Life Laboratory, Stockholm, Sweden). The resulting reads were mapped and variants called, filtering for variants in the coding regions and excluding indels, using the Partek Flow lab edition software and DNA-Seq Toolkit for Partek Flow.

### High-throughput drug screen for mono-therapy

Cells were seeded at the cell densities indicated in Supplementary Table 1B in 384-well plates upon different drug treatment for 72 hours on black clear bottom imaging quality 384-well plates (Corning, cat # 3985). Prior to fixation, cells were incubated with 5 µM EdU reagent (Thermo Scientific) for 30 min. Then cells were fixed with 4% PFA, permeabilized with 0.3% Triton-X100, blocked with 2% BSA. EdU detection was performed first according to the manufacturer’s protocol. For other biomarkers, cells were incubated with phalloidin (Thermo Scientific) or different primary antibodies (see Supplementary Materials and Methods) at 4 °C overnight before counterstaining with DAPI (Thermo Scientific). Imaging was performed with Olympus scan^R high-content imager with a 10x objective. Intensities per field were collected and after normalizing to DMSO, median values (%) were displayed in heatmaps, which was generated via Morpheus (https://software.broadinstitute.org/morpheus).

Drugs were dissolved in DMSO at indicated stock concentration in Suppl. Table 1A, and stored in −20 °C.

### Cell cycle analysis

Cell cycle analysis was performed as described in (35) using 5-Ethynyl-2′-Deoxyuridine (EdU) Staining and Flow Cytometry. In brief, prior to harvesting, the cells were pulsed with 10 µM EdU for 90 min, harvested, stained according to manufacturer’s instructions, and analyzed by FACSCalibur (BD Biosciences). Cell cycle analysis was performed by FlowJo software.

### Senescence-associated β-galactosidase activity (SA-β-gal) assays

Drug induced senescence was investigated by assessing β-galactosidase activity using the quantitative SA-β-gal MUG assay (36) or by x-gal senescence-associated β-galactosidase staining, pH 6 (#9860 staining kit, Cell Signaling Technology), as described in detail (37) (see also Supplementary Materials and Methods). Cells were treated with indicated drugs for 3-5 days before harvesting. All MUG assay reagents were purchased from Sigma-Aldrich. The amount of fluorescent product was measured using a Tecan microplate reader (Tecan Trading AG). X-gal SA-β-gal staining was assessed by light microscopy.

### RNA and protein extraction, real-time PCR and western blot analysis

Total RNA was extracted using the Aurum™ Total RNA Mini Kit (Bio-Rad) following the standard manufactureŕs protocol. For relative quantification of mRNAs, one µg total RNA was reverse transcribed to cDNA using the iScript™ cDNA Synthesis Kit (Bio-Rad) according to the manufacturer’s instructions. Real-time PCR was conducted with SsoAdvanced™ Universal SYBR® Green Supermix in 384 well format using CFX384 real-time system (Bio-Rad). For primer sequences, see Supplementary Materials and Methods. The relative expression of genes was determined using the Bio-Rad CFX Maestro software. The data were presented as 2(-ΔΔCT). Protein extraction was performed using RIPA buffer (25 mM Tris-HCl pH 7.6, 150 mM NaCl, 1% NP-40, 1% sodium deoxycholate, 0.1% SDS), 1 mM NaOV, protease and phosphatase inhibitors as described (38). Protein concentration was measured using BCA kit according to manufacturer’s protocol (Thermo Scientific).

40-50 µg of protein was loaded on a 4-12% NuPage Bis-Tris gel (Thermo Scientific), transferred onto a 0.45 µm PVDF membrane (Millipore), blocked using 5% BSA/milk and incubated with the primary overnight at 4°C. Following day, the membranes were washed with 1X TBST (3X, 10 min wash each) and incubated with the appropriate HRP-labeled secondary antibodies (Cell signaling Technology) for 1h, followed by ECL based detection using Image Quant LAS 4000 (GE Healthcare Europe GmbH, Freiburg, Germany). Primary antibodies used are listed in Supplementary Materials and Methods.

### High-throughput drug screen for combination-therapy

Approximately 3000-5000 cells were seeded in 96-well plates and treated with a variety of drug combinations for 72 hours. Cells were fixed with 4% paraformaldehyde (Sigma), permeabilized with 0.5% Triton X-100 (Sigma), blocked with 4% BSA (Sigma), and then stained with phalloidin (Thermo Scientific) and counterstained with DAPI. Cell number, nuclear/cell size and EdU incorporation were examined by automated fluorescent imaging system ImageXpress Micro (Molecular Devices). Image analysis were done via CellProfiler and mean values were presented as heatmaps via Morpheus (https://software.broadinstitute.org/morpheus).Combenefit software was used to calculate combination index (39), where HSA model was applied for synergism/antagonism definition and displayed as heatmaps via online Morpheus software (https://software.broadinstitute.org/morpheus/).

### Analysis of immune markers on human melanoma cells

Treated tumor cells were harvested and washed in PBS for 15 min at room temperature. Cells were incubated with antibodies diluted in FACS buffer (PBS + 1% BSA) for 30 min at 4°C. Flow cytometry was performed using a Novocyte 2000 flow cytometer (Acea Biosciences, San Diego, CA, USA). Primary antibodies used are listed in Supplementary Materials and Methods. LIVE/DEAD Fixable Aqua Dead Cell Stain (Thermo Scientific) was included as a viability dye in all experiments to enable exclusion of non-viable cells in our analyses. Analysis was done using FlowJo software version 10 (Treestar). The results are expressed as geomean fluorescence intensity (geoMFI) of at least three independent experiments.

### Melanoma mouse model

All animal experiments were performed in accordance with the guidelines of the Karolinska Institutet and approved by the Stockholm North Ethics Committee for Animal Research (ethical approval 10025/23). Five- to six-week-old female C57BL/6J mice were subcutaneously injected with 2.5 × 10^5^ YUMM1.7 cells mixed in 100uL PBS/Geltrex (75:25). Tumor size was measured and recorded three times per week using the formula V = (L x W^2^)/2. Drug treatment began when the tumors were palpable. Animals were divided into treatment groups and treated with 180uL oral gavage according to these groups: Group 1: vehicle (0.5% methylcellulose, 0.5% Tween80), Group 2: 80mg/kg palbociclib diluted in vehicle, Group 3: 60mg/kg crizotinib diluted in vehicle, Group 4: 80mg/kg palbociclib + 60mg/crizotinib diluted in vehicle. At the end of treatment, tumors were harvested, weighed, and subjected to flow cytometric analysis. To deplete CD8+ T cells, 200ug/mouse InVivoMab anti-mouse CD8a clone 2.43 (Bioxcell, BR0061) or InVivoMab rat IgG2b isotype control (Bioxcell, BE0090) were diluted in InVivoPure pH7 dilution buffer (Bioxcell, IP0070), and injected i.p. two days before the start of drug treatment, followed by administration of 100ug/mouse every 4 days during the course of tumor growth. The depletion efficiency was above 98 % as determined by flow cytometry.

### Immune infiltration in YUMM1.7 tumors

Harvested tumors were cut into small pieces in RPMI 1640 media serum free using razor blades. Enzymatic and mechanical dissociation was performed using the mouse tumor dissociation kit (130-096-730, Milteniy) and the gentleMACS™ Dissociator (130-093-235, Milteniy) according to the manufacturer’s specifications. Cell debris was removed by centrifugation 100xg for 10 minutes. Red blood cells were removed using RBC lysis buffer (MIK1512, Karolinska Hospital) for 5 minutes at room temperature. FcγR was blocked before staining using Fc Block anti mouse CD16/CD32 (clone 93, Biolegend) for 5 minutes at 4°C. The primary antibodies used for flow cytometry are listed in Supplementary Materials and Methods. All data were acquired on a BD LSRII or Fortessa flow cytometer (BD Bioscience). Analysis was done using FlowJo software version 10 (Treestar). The quality control of the raw data was performed using the flowAI plugin in FlowJo and only the “GoodEvents” population was kept for further analysis (40).

### Statistical analysis

One-way ANOVA with post-hoc Dunnett’s multiple comparison test was applied used and performed via GraphPad Prism version 10.00 (for Windows, GraphPad Software, La Jolla California USA, www.graphpad.com), at a level of significance α= 0.05.

### Data Availability

The data generated in this study is available upon request from the corresponding author.

## Results

### Drug therapy-induced senescence screening in melanoma cell lines

To identify drugs with senescence-inducing activity in melanoma, a senescence screen was set up using a panel of eleven human melanoma cell lines (Fig. 1A and Table 1), representing some of the most common driver mutations in melanoma. A set of conventional and targeted drugs and compounds of relevance for melanoma and pro-senescence therapy, either already in clinical use or at an experimental stage were selected, including vemurafenib (4), trametinib (5), BKM120 (41), palbociclib (42), crizotinib (43), and seliciclib (44), inhibiting BRAF^V600E^, MEK1/2, PI3K, CDK4/6, ALK/MET/ROS1, CDK2/7/9, respectively, and the DNA alkylating agent temozolomide (45). In addition, the p53-activating molecules Nutlin-3A (46), RITA (47) and APR-246 (Eprenetapopt) (48) were included (Table 1). The screening was performed using a high-throughput multicolor immunofluorescence microscopy system (Fig. 1A, Suppl. Tables 1 and 2), staining for DAPI (cell number), phalloidin (cell size), EdU incorporation (replication), H3K9me3 (senescence-associated heterochromatin foci) and p53, thereby covering many essential markers of senescence suitable for fluorescence staining (15) as well as HLA class I (antigen recognition by T cells). The criterion for a major senescence response was the combination of reduced cell number and EdU incorporation, increased cell size, elevated H3K9me3 and p53 expression. While cell cycle arrest is an obligatory criterion, increased expression of only one or some other markers would be considered “partial senescence” or a “senescence-like” phenotype.

**Figure 1.**
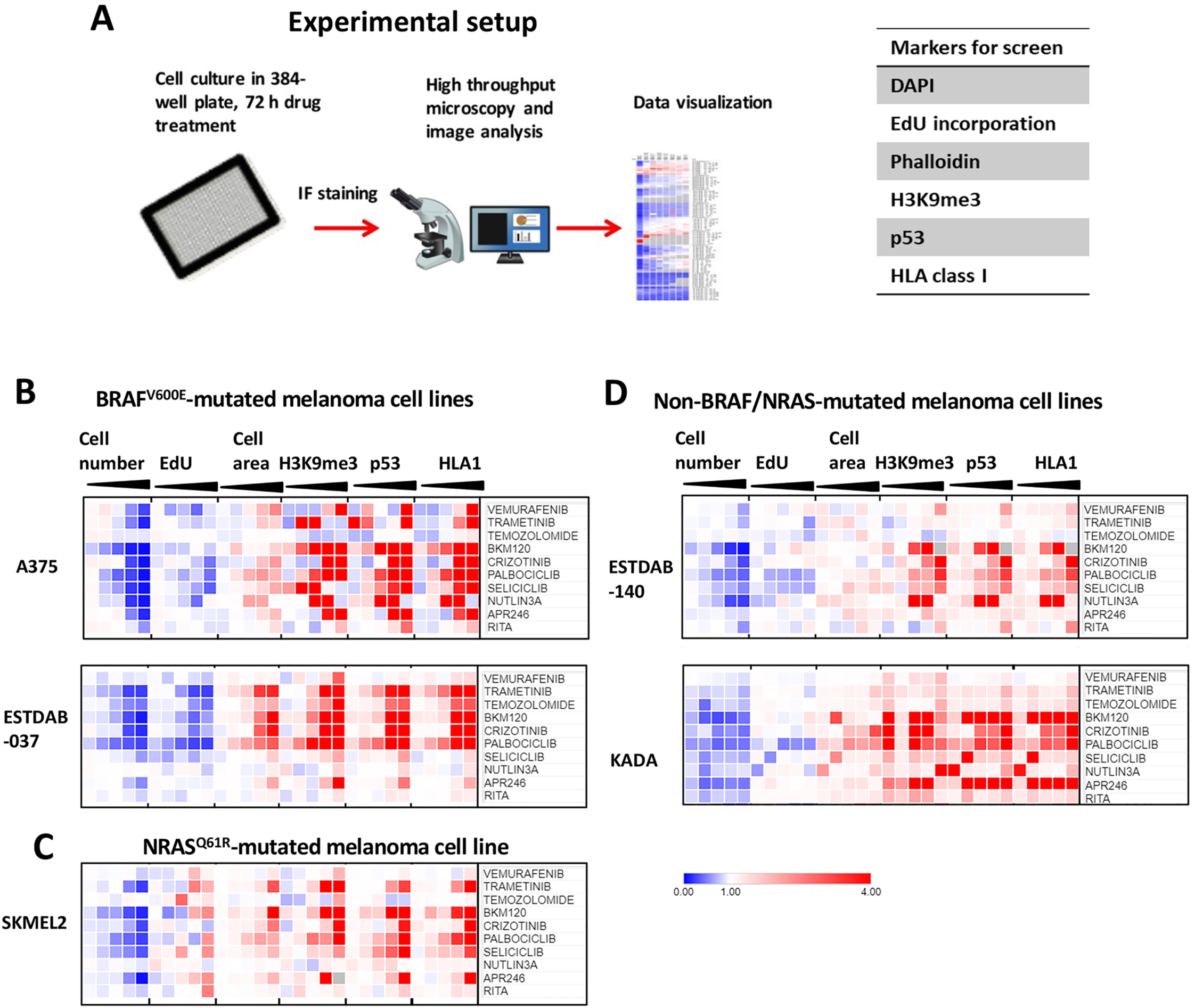
Screen for drug-induced senescence in human melanoma cell lines (A) Experimental set-up and workflow of the drug-induced senescence screen. Right panel: Markers used in the screen, DAPI (cell number), EdU incorporation, phalloidin (cell area), H3K9me3, p53 and HLA class I immunofluorescence staining. (B-D) Heatmaps showing data from measurements of the six biomarkers in response to 11 selected drugs after 72 hours of treatment in 5 selected melanoma cell lines. Drug concentrations are specified in Supplementary Table 1A, and increasing concentration are illustrated as wedges. (B) A375 and ESTDAB037, both with BRAF^V600E^ mutation; (C) SKMEL2 with NRAS^Q61R^ mutation; (D) ESTDAB140 and KADA, with non-BRAF/NRAS mutations. Intensity per field was normalized to DMSO, and median values (%) were presented here with indicated color code.

The results of the screen are presented as heatmaps in Fig. 1B-D (five selected representative cell lines) and Suppl. Fig. S1A-C (all 11 cell lines). Inhibition of BRAF^V600E^ with vemurafenib induced a major senescence response in two of the *BRAF^V600E^*-mutant cell lines, A375 and ESTDAB-049, but not in SKMEL28 and ESTDAB37 (Fig. 1B and Suppl. Fig. S1A), indicating that the latter possess an intrinsic resistance to BRAF^V600E^ inhibition, possibly due to additional driver mutations and/or copy number alterations (Table 1). We observed a clear senescence response to the MEK1-inhibitor trametinib in all *BRAF^V600E^*-mutant cells lines, but poor response in two out of three cell lines with *NRAS* mutation and in the non-*BRAF/NRAS* mutant cell lines, (Fig. 1C and Suppl. Fig. S1B, C). Most cell lines were essentially non-responsive to temozolomide at the concentrations used.

Interestingly, palbociclib, BKM120 and crizotinib efficiently induced senescence in all or most cell lines irrespective of *BRAF*/*NRAS* mutation status. Likewise, the CDK2 inhibitor seliciclib induced senescence in many of the cell lines of all categories. The p53:MDM2 interaction inhibitors Nutlin-3A and RITA induced senescence responses to a greater extent in wt *TP53* cell lines and less in cell lines with mutant *TP53* (Table 1) and vice versa for APR-246, which targets mutant p53 and disrupts redox homeostasis (49).

### Palbociclib, BKM120 and crizotinib induce senescence and HLA class I expression in melanoma cells irrespective of *BRAF*/*NRAS* mutation status

The results from the senescence screen suggested that palbociclib, BKM120 and crizotinib induced either prominent or partial senescence in all eleven melanoma cell lines regardless of *BRAF*/*NRAS* mutation status, including *BRAF*-mutant cells resistant to vemurafenib and trametinib. The senescence response to these drugs was further validated in five selected representative cell lines; vemurafenib sensitive/resistant *BRAF^V600E^* mutant (A375, ESTDAB37), *NRAS^Q61R^* mutatant (SKMEL2), non-*BRAF*/*NRAS*-mutant (ESTDAB140, KADA).

The results confirmed that vemurafenib induced senescence in a dose-dependent manner in A375 cells, as evidenced by an increase in cell/nuclear size and H3K9me3 staining intensity along with reduced EdU incorporation and an accumulation of cells in the G1 or G2 phases of the cell cycle (Fig. 2A-C, Suppl. Fig. S2). In contrast, vemurafenib failed to induce most markers of senescence in the *BRAF^V600E^*-mutant ESTDAB37 cells and in the non-*BRAF*-mutant cell lines. Trametinib induced a good senescence response in *BRAF*/*NRAS*-mutant but a weak response in the non-*BRAF*/*NRAS*-mutant cells (Fig. 2A-C, Suppl. Fig. S2). Treatment with the CDK4/6 inhibitor palbociclib significantly increased cell/nuclear size and H3K9me3, decreased EdU incorporation and caused G1 arrest, indicative of major senescence response in all five lines, but did not affect cell size in ESTDAB140 cells (Fig. 2A-C, Suppl. Fig. S2). The PI3 kinase inhibitor BKM120 and the MET/ALK/ROS1 inhibitor crizotinib induced significant changes for most senescence markers in all the cell lines, although with some differences between cell lines and senescence markers (Fig. 2A-C, Suppl. Fig. S2).

**Figure 2.**
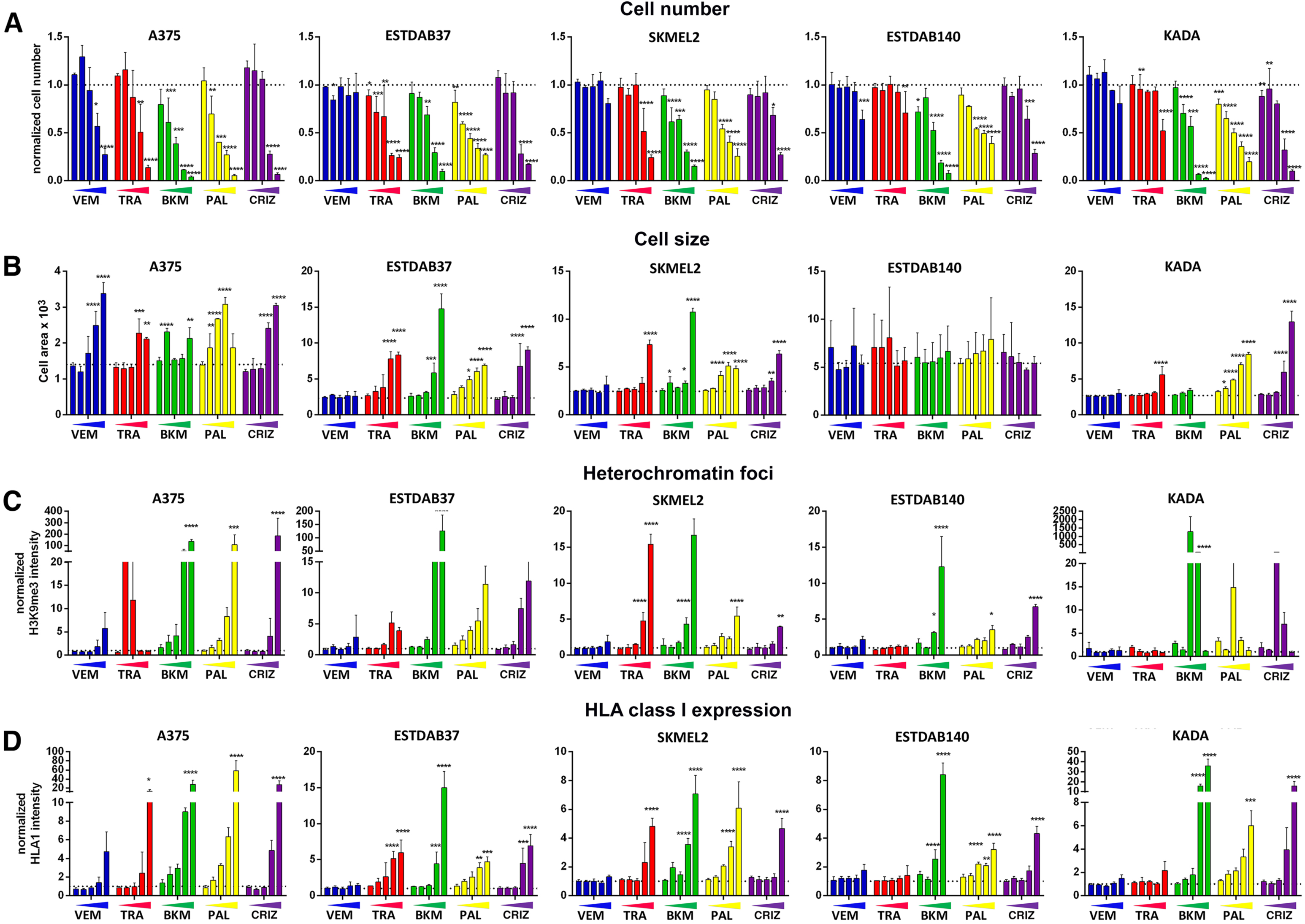
Validation of senescence induction in response to selected drugs in selected melanoma cell lines. Single-drug treatment of A375, ESTDAB37, SKMEL2, ESTDAB140 and KADA with vemurafenib (VEM), trametinib (TRA), BKM120 (BKM), palbociclib (PAL) or crizotinib (CRIZ) for 3 days. Bar charts illustrating cell number (A), cell size (B), intensity of H3K9me3 (C) and HLA class I (D) stainings. Drug concentrations are specified in Supplementary Table 1A, and increasing concentration are illustrated as wedges. Data was normalized to DMSO treatment and mean±SEM values are presented. The value for DMSO for each marker is indicated as a dotted line in the figure. Blue: VEM; red: TRA; green: BKM; yellow: PAL; purple, CRIZ. * p< 0.05; ** p< 0.01, *** p<0.001, **** p<0.0001.

We also validated whether senescence-inducing drugs affected the expression of HLA class I (Fig 1, Suppl. Fig. 1, Fig 2 D). Vemurafenib increased HLA class I expression in A375 cells, but not in ESTDAB37 cells or any of the other cell lines. Trametinib increased HLA class I expression in the BRAF or NRAS-mutant cell lines and slightly in KADA cells, which has a truncating mutation in *NF1* – a regulator of the RAS/MAPK signaling pathway. Remarkably, palbociclib, BKM120 and crizotinib strongly induced expression of HLA class I in all the cell lines.

In summary, while vemurafenib and trametinib induced senescence and HLA class I expression in a subset of cell lines related to *BRAF*/*NRAS* mutation status, palbociclib, BKM120 and crizotinib treatments resulted in a pronounced senescence response and HLA class I expression in all cell lines regardless of *BRAF*/*NRAS* mutation status.

### Palbociclib, crizotinib and BKM120 synergize with vemurafenib to induce senescence in vemurafenib-resistant *BRAF^V600E^*-mutant melanoma cell lines

The results of the mono-treatments showed that palbociclib, BKM120 and crizotinib induced senescence in all four BRAF^V600E^-mutant cell lines, including ESTDAB37 cells that were resistant to vemurafenib-induced senescence. We were therefore interested in whether these drugs would synergize with vemurafenib to overcome vemurafenib resistance. We selected A375 as a control for vemurafenib-sensitive cells, while A375-VR4 (a vemurafenib-resistant subline derived from A375) (33) and ESTDAB37 served as two vemurafenib-resistant cell lines with BRAF^V600E^ mutation.

As shown in the heatmaps in Fig. 3, the combination of vemurafenib and palbociclib efficiently reduced cell number and induced cell cycle arrest and cell enlargement, i.e. signs of senescence induction in A375, A375-VR4 and ESTDAB37 cells. Calculation of the synergy between the two drugs based on the data of cell size using combination index showed a strong synergistic effect as visualized in the heat maps to the far right on each panel (Fig. 3A-C). The synergism was particularly strong in the vemurafenib-resistant lines A375-VR4 and ESTDAB37. The combination of vemurafenib with temozolomide, BKM120, crizotinib or trametinib had some synergistic effects in all three cell lines, but less than the combination vemurafenib + palbociclib (Fig. 3, Suppl. Fig. S3A).

**Figure 3.**
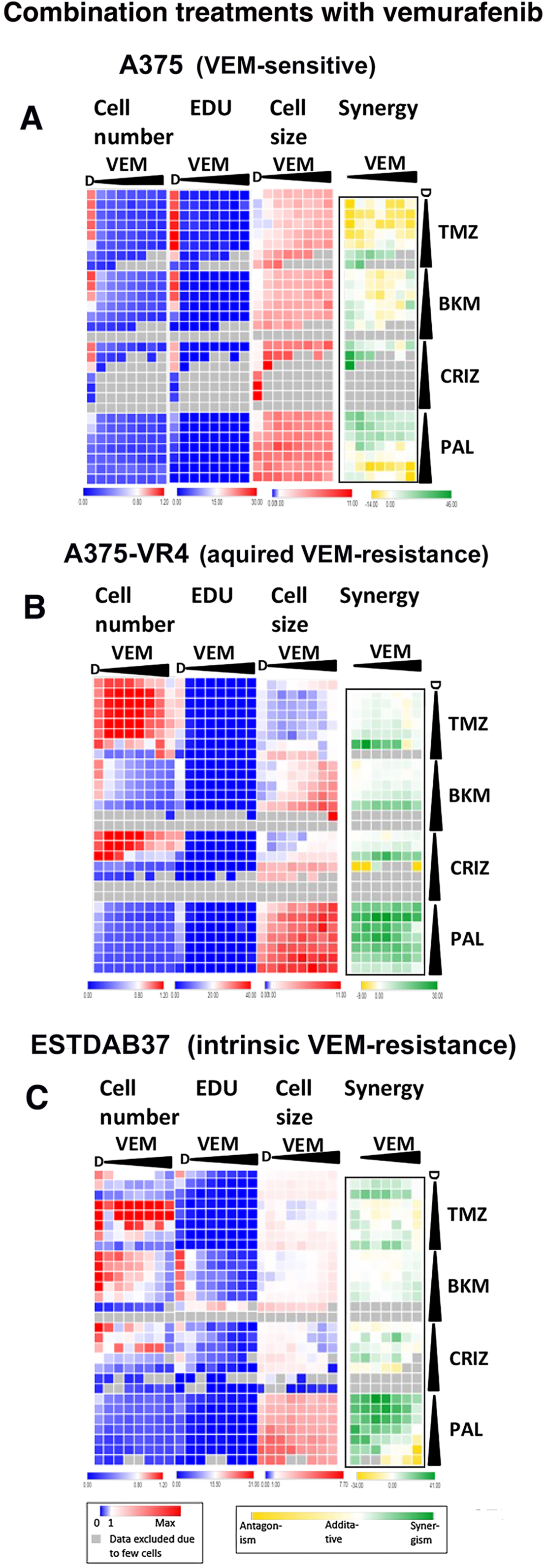
Drug combination senescence screen using vemurafenib-sensitive and vemurafenib-resistant melanoma cell lines. Combination treatment using vemurafenib (VEM) together with temozolomide (TMZ), BKM120 (BKM), crizotinib (CRIZ) or palbociclib (PAL) at 2-fold serial dilutions for 72 hours in A) A375, B) A375-VR4 and C) ESTDAB37 cells. The highest concentrations of the drugs in each combination are listed in Suppl. Table 2A and increasing concentrations for each drug are illustrated as black wedges. D=DMSO. The heatmaps show data representing cell number, percentage of EdU positive cells, cell size and combination synergy index based on cell size in A375, A375-VR4 (acquired VEM-resistance) and ESTDAB37 (intrinsic VEM-resistance) cells. Cell number and cell size were normalized to DMSO-treated cells. Missing data due to too low cell number as a result of cell death is indicated as grey. Green = synergism, yellow = antagonism.

We also investigated combination treatments with the MEK1/2 inhibitor trametinib together with palbociclib, BKM120, crizotinib, temozolomide, APR-246 or RITA in the non-*BRAF*/*NRAS* mutant cell line KADA, which has a truncating mutation in *NF1* – a regulator of the RAS pathway. Suppl. Fig. S3B shows that all these drugs and compounds synergized with trametinib to some extent.

### Palbociclib and crizotinib synergistically enhance senescence in melanoma cells irrespective of *BRAF*/*NRAS* mutation status

Since palbociclib, BKM120 and crizotinib induced prominent or partial senescence in all melanoma cell lines irrespective of *BRAF*/*NRAS* mutation status (Fig. 1, 2 and Suppl. Fig. S1, S2), we next addressed the question whether these three drugs might work cooperatively. Therefore, we performed a screen with the same layout as above with the combination palbociclib+BKM120, palbociclib + crizotinib and BKM120 + crizotinib on A375, A375-VR4, ESTDAB 37, KADA and ESTDAB140 cells. While the combinations palbociclib+BKM120 and BKM120+crizotinib produced rather weak synergies except for palbociclib + BKM120 in KADA cells (Fig. 4A, Suppl. Fig. S4A, D), the combination palbociclib plus crizotinib showed strong synergistic effects on senescence induction in all five melanoma cell lines (Fig. 4A and Suppl. Fig. S4A). Considering the limited clinical activity and tolerability of BKM120 in recent clinical trials (50), we therefore chose to focus on the palbociclib/crizotinib combination in the subsequent experiments.

**Figure 4.**
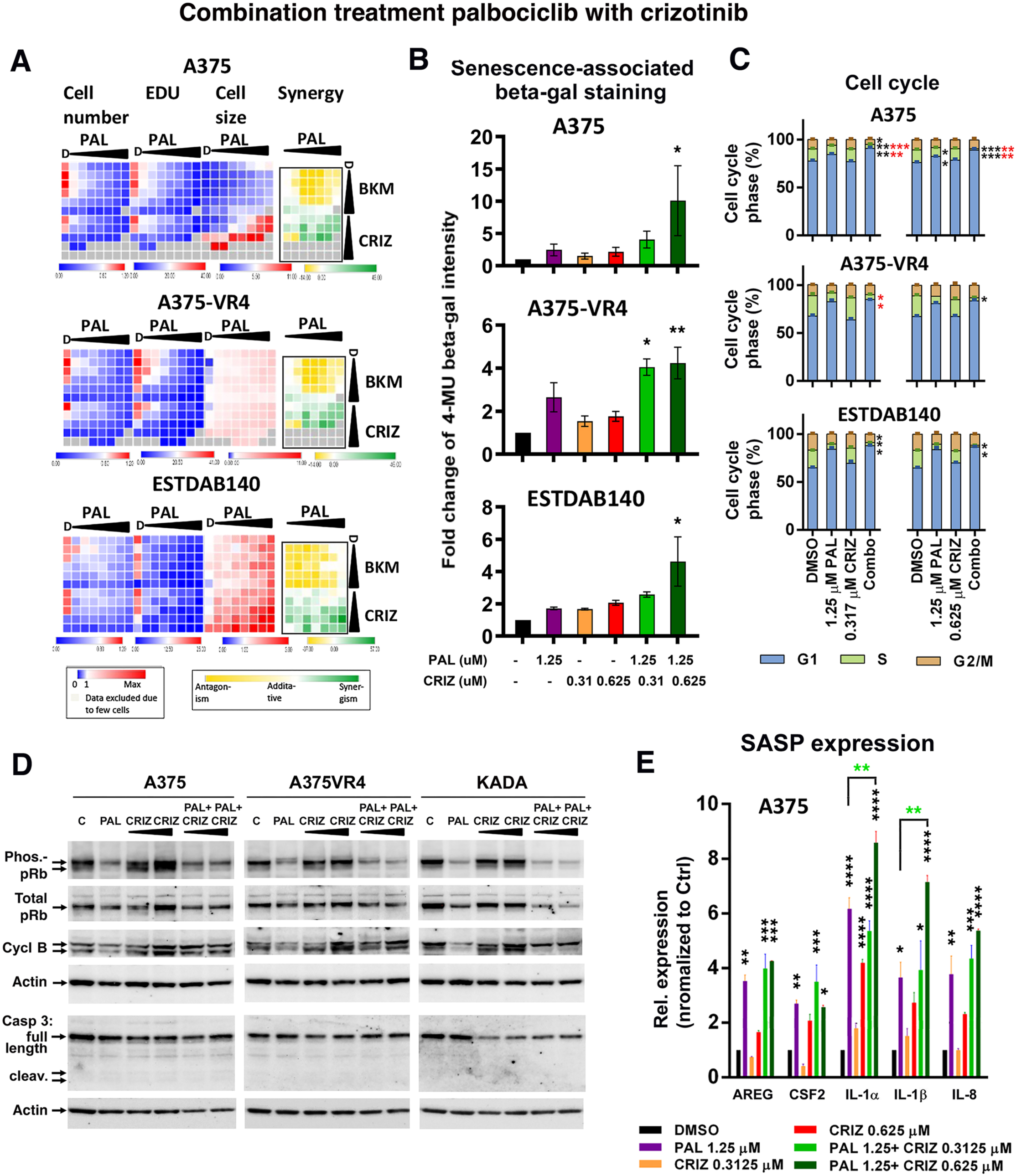
Drug combination senescence screen using selected melanoma cell lines. (A) Combination treatment using PAL together with BKM or CRIZ at 2-fold serial dilutions for 72 hours. The highest doses of the drugs are listed in Suppl. Table 2B. The heatmaps show data representing cell number, percentage of EdU positive cells, cell size and combination synergy index based on cell size in A375, A375-VR4, and ESTDAB140 cells. Missing data due to too low cell number as a result of cell death is indicated as grey. Green = synergism, yellow = antagonism. (B) Fluorescent β-galactosidase assay (MUG) after mono or combination treatment with PAL and CRIZ as indicated. (C) Cell cycle analysis after mono or combination treatment with PAL and CRIZ in indicated cell lines for 72 hours. (D) Western blot analysis of S780 phosphorylated pRb, total pRb, cyclin B1 and cleaved caspase 3 after mono or combination treatment with a fixed concentration of PAL (1.25 μM) and increasing concentrations of CRIZ (0.31 and 0.63 μM as illustrated by wedges) in the indicated cell lines for 72 hours. (E) Relative fold change of SASP-related gene expression in A375 cells in response to combination treatment using palbociclib (PAL) and crizotinib (CRIZ) for 72 hours. Biological triplicates were presented as mean±SEM and one-way ANOVA was applied to compare drug treatment versus mono-therapy, * p< 0.05; ** p< 0.01, * p< 0.05; ** p< 0.01, *** p<0.001. Black stars represent significance DMSO control vs. treatments and are placed above columns in the graphs in (B) and (E), red and green stars represent significance CRIZ vs. combination and PAL vs. combination treatments, respectively, and are placed above brackets between columns in (E).

We first analyzed SA-β-gal activity, a well-known senescence marker. While palbociclib and crizotinib alone increased SA-β-gal activity as expected (Fig. 4B and Suppl. Fig. S4B), combined treatment with palbociclib and crizotinib significantly enhanced SA-β-gal activity in all five cell lines. Further, we observed a significantly increased percentage of cells in the G1 or G2/M phase and decreased number of cells in S phase of the cell cycle upon combined compared with mono treatments in all cell lines (Fig. 4C and Suppl. Fig. S4C). We also examined additional senescence markers, phosphorylation of the retinoblastoma protein (pRb) and the expression of cyclin B1 in three of the cell lines: A375, A375-VR4, and KADA. As expected, treatment with palbociclib alone resulted in decreased pRb phosphorylation (i.e. pRb activation) and reduced cyclin B1 expression in all three cell lines, while crizotinib treatment alone did not affect either marker. Combined treatment with palbociclib and crizotinib did not, however, lead to further reduction in pRb phosphorylation or cyclin B expression compared with palbociclib alone (Fig. 4D). This is likely because palbociclib directly controls pRb phosphorylation through inhibition of CDK4/6. Analysis of full-length and cleaved caspase 3 showed no evidence of apoptosis induction by palbociclib alone, crizotinib alone, or their combination at the drug concentrations used (Fig. 4D, second panel from the bottom). Taken together, these results demonstrate that the combination of palbociclib and crizotinib synergistically enhances the senescence response with respect to most markers of senescence in melanoma cells.

### Treatment with palbociclib and crizotinib modulates the expression of SASP factors and immune-related receptors on melanoma cells

We next monitored the expression of eight reported SASP-related genes (15) in the five selected melanoma cell lines upon mono- or combination-treatments with palbociclib and crizotinib. The expression of the five SASP genes, IL-1α, IL-1β, AREG, IL-8 (CXCL8), and CSF2, increased signifcantly after combination treatment in all (IL-1α and IL-1β) or 4 out of 5 cell lines (AREG, CSF2 and IL-8) as shown in Fig. 4E and Suppl. Fig. 5. Mono-treatment with palbociclib or crizotinib also significantly increased the expression of these genes in the majority of the cell lines, except for IL-1β and CSF2 after crizotinib treatment, which were induced only in one and two of the cell lines, respectively. Compared with palbociclib treatment alone, the combination treatment significantly enhanced the expression IL-1α and IL-1βin A375 cells, while in A375-VR4 cells the expression IL-1β and IL-8 was somewhat reduced relative to DMSO, Fig. 4E and Suppl. Fig. 5B). The expression of the other three SASP-related genes (CXCL1, IL-6 and MMP3) increased significantly in only 1-3 of the cell lines after the mono- or combination treatments (Suppl. Fig. 5). In general, palbociclib was more efficient inducing the SASP genes in A375 and A375-VR4 cells (although the amplitude was higher in the former), while crizotinib was more efficient in KADA and ESTDAB37 cells. In KADA cells, the inductions of many of the SASP genes were more efficient with crizotinib alone compared with the combination treatment (Suppl. Fig S5E).

Taken together, this suggests that the most consistent SASP response, in particular with respect to IL-1α, IL-1β, AREG, IL-8, and CSF2, was observed after combination treatment with palbociclib and crizotinib, but also mono--treatments with palbociclib or crizotinib were often efficient and in some cases more powerful than the combination of the two drugs.

We next investigated the expression of immune receptors of relevance for natural killer (NK) and T-cell recognition and killing of A375, a long-term established cell line, A375-VR4, it’s VEM-resistant variant and KADA, a recently established melanoma cell line by FACS. In all three cell lines, palbociclib treatment alone significantly increased the expression of HLA class I, while crizotinib treatment did not affect HLA class I expression to a major extent. Combination of the two drugs lead to a further significant upregulation compared with palbociclib alone in KADA, but not in A375 or A375-VR4 cells (Fig. 5A and Suppl. Fig. S6A). Expression of HLA class II was upregulated in all cell lines by palbociclib alone and by crizotinib alone in KADA cells, and a significant further upregulation after combination treatment in KADA cells (Fig. 5B and Suppl. Fig. S6B). Expression of PD-L1, which inhibits T and NK cell activation, increased significantly by mono-treatment with palbociclib in KADA cells, and a trend towards an increase in the other two cell lines, as well as after crizotinib mono-treatment in KADA cells. The combined treatment further increased PD-L1 expression significantly in A375 but not in the other two cell lines (Fig. 5C and Suppl. Fig. S6C).

**Figure 5.**
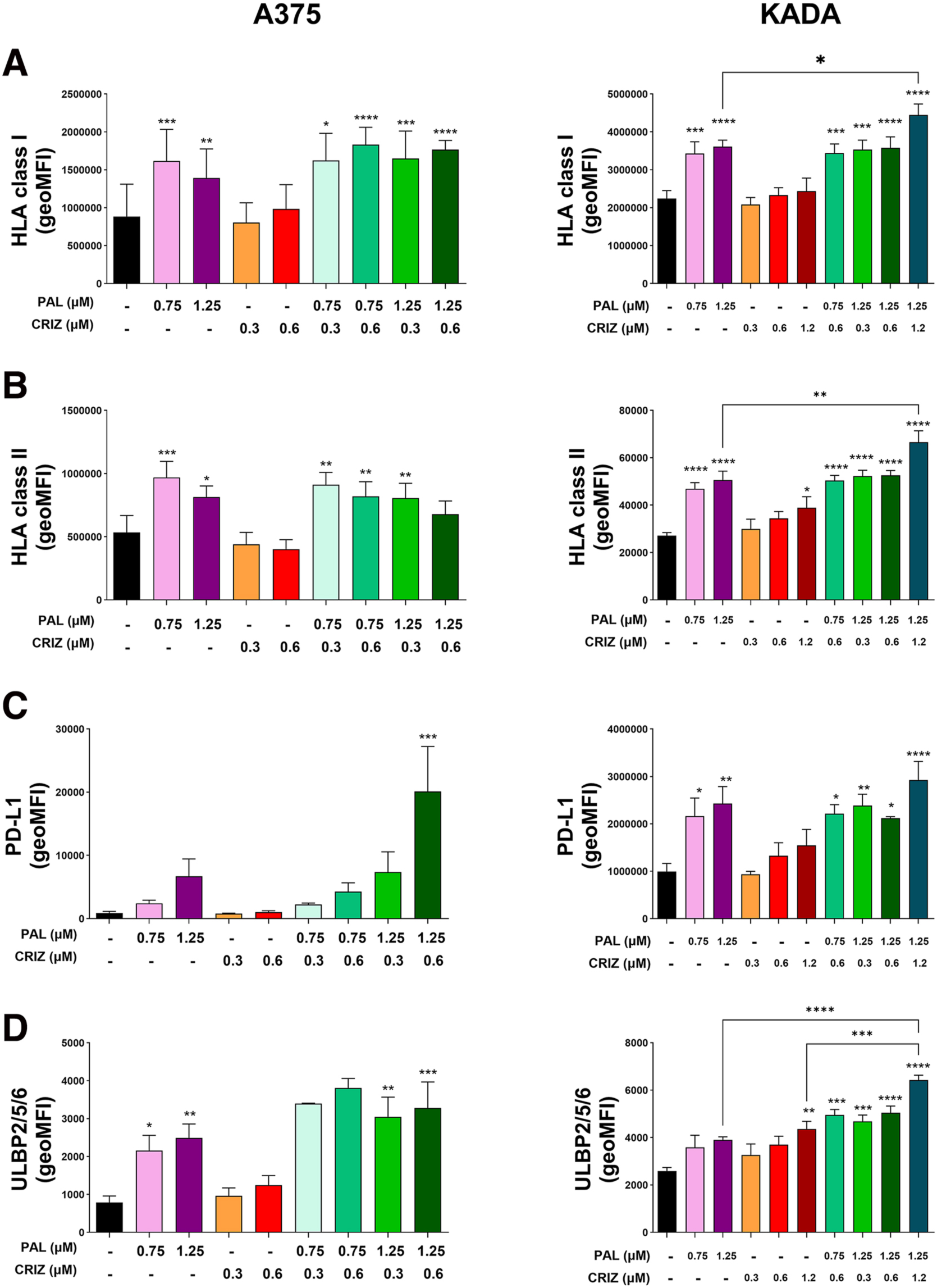
Expression of immune markers regulating recognition by T cells and NK cells upon combination treatment with palbociclib and crizotinib. A375 (left panel) and KADA (right panel) cells were treated with indicated concentrations of PAL and/or CRIZ for 72 hrs followed by analysis of (A) HLA class I, (B) HLA class II, (C) PD-L1 and (D) ULBP2/5/6 by flow cytometry using specific antibodies. * p< 0.05; ** p< 0.01, *** p<0.001, **** p<0.0001. Stars placed above columns in the graphs represent significance DMSO control vs. treatments, and stars placed above brackets between columns represent significance mono-treatment vs. combination treatments.

Further, palbociclib mono-treatment upregulated the expression of ULBP 2/5/6, which is a ligand of NKG2D, a receptor involved in NK and T cell activation, in A375 and A375-VR4 and to some extent in KADA cells, while crizotinib treatment alone only increased its expression in KADA cells (Fig. 5D and Suppl. Fig. S6D). The combination treatment led to further <u>significantly</u> increased expression compared with the mono-treatments in KADA, but not in the other two cell lines. In KADA cells we also investigated the expression of MIC A/B, another NKG2D ligand, which was significantly induced by palbociclib, but not by crizotinib treatment alone, while the combination treatment did not lead to any further increase (Suppl. Fig. 6E). However, the combination of palbociclib and crizotinib significantly increased the expression of DR5 but not of FAS (both involved in killing by NK and T cells) and a trend towards an increase in DR5 expression was observed after palbociclib mono-treatment (Suppl. Fig. S6F, G).

Taken together, this indicates that molecules involved in recognition both by T and NK cells are generally upregulated by palbociclib, some also by crizotinib, and in some cases combination treatment led to a further increased expression of these molecules. On the other hand, the immune checkpoint molecule PD-L1 was also upregulated by palbociclib and in some cases further increased by the combination treatment.

### Combination treatment with palbociclib and crizotinib inhibits melanoma tumor growth *in vivo* and increases infiltration of CD8+ T-cells and M1-like macrophages into tumor tissue

To address whether senescence induced by palbociclib and crizotinib would affect melanoma tumor growth *in vivo* and to assess any involvement of immune cells in such a response, we next utilized the mouse melanoma cell line YUMM1.7 for syngeneic transplantation into immunocompetent mice. YUMM1.7 is derived from a mouse melanoma tumor carrying BRAF^V600E^ mutation and deletions of PTEN and CDKN2A (34), resembling a subset of human melanomas. First, we studied the senescence response of YUMM1.7 cells *in vitro*. Combination treatment with palbociclib and crizotinib significantly reduced cell growth compared with DMSO and with mono-treatments (Suppl. Fig. 7A). Further, palbociclib and to a lesser extent crizotinib treatment alone led to increased β-gal staining and cell size, indicative of senescence induction, which was further increased by the combination treatment (Suppl. Fig. 7B).

To validate the effect of palbociclib and crizotinib combination treatment on melanoma growth *in vivo*, YUMM1.7 cells were transplanted into the flanks of immunocompetent C57BL/6J mice. Upon palpable tumors, the mice were treated with palbociclib and/or crizotinib or with vehicle for 2 weeks. While single treatments had some effect on tumor growth, the combination treatment significantly inhibited tumor growth and reduced tumor volume (Fig. 6A, B), but none of the treatments had any significant effect on mouse weight (Suppl. Fig. S7C), suggesting lack of serious side effects of the treatments.

**Figure 6.**
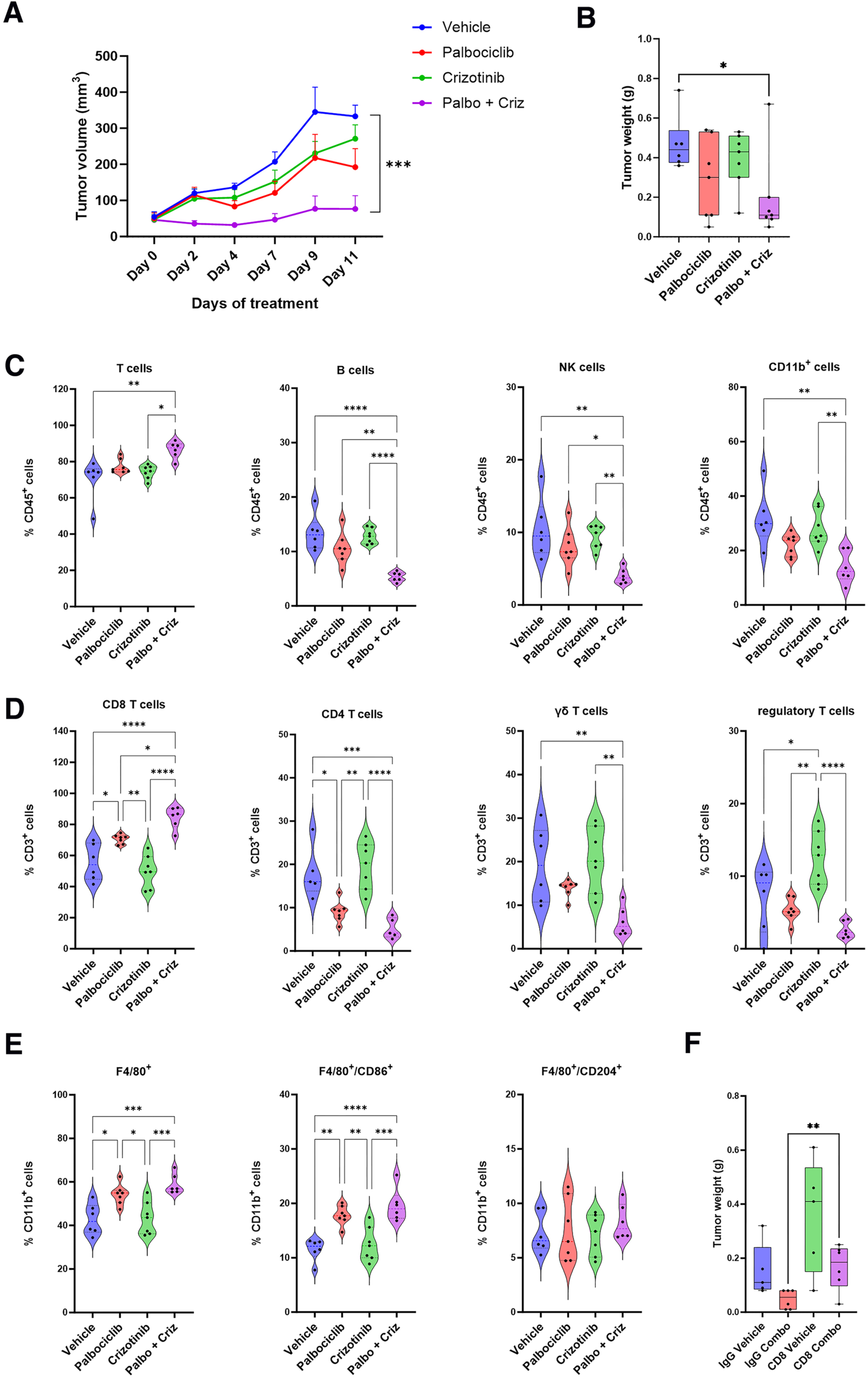
Effects of combination treatment using palbociclib (Palbo) and crizotinib (Criz) on tumor growth and immune cell infiltration after syngeneic transplantation of YUMM1.7 melanoma cells into immunocompetent C57BL6 mice. (A) YUMM1.7 tumor volume measurement during treatment period (11 days), data shown are Mean ±SEM (B) Violin box showing YUMM1.7 tumor weight measurement at endpoint (C-E) Violin box showing frequencies of infiltrating immune cells as determined by flow cytometry. (C) Analysis of total T, B, NK, and myeloid cells. (D) Analysis of subpopulations of T cells as indicated (D) Analysis of total, M1-like, and M2-like macrophages. Differences between vehicle and treatment conditions were assessed by one-way ANOVA test, * p< 0.05; ** p< 0.01; ***p<0.001, ****p<0.0001 - Represented points for (A), (B) and (C-E): Vehicle, n=6; Palbo, n=7; Criz, n=7; Palbo + Criz, n=7. (F) Effects of CD8+ T cell depletion on tumor growth. Anti-mouse CD8a or IgG2b isotype control antibodies were injected i.p. into tumor-bearing mice two days before the start of drug treatment, followed by administration every 4 days during the course of the experiment. Difference between IgG Combo and CD8 Combo was assessed by unpaired t-test, ** p= 0.0096. Represented points for (F): IgG Vehicle, n=5; IgG Combo, n=6; CD8 Vehicle, n=5; CD8 Combo, n=6.

We then assessed the effect of the treatments on the immune cell profile in the tumor microenvironment by analyzing different immune cell populations (including T, B, and NK cells, macrophages and neutrophils as well as subtypes of T cells and macrophages) from dissociated tumors by flow cytometry. The combination treatment had in general a more pronounced effect on immune cell composition than either monotherapy or vehicle (Fig 6C-E). We observed a significant enrichment of T cells and significant reductions of B cells, NK cells and myeloid cells after combination treatment as compared to the mono-treatments and vehicle (Fig. 6C). Among T cells, the combination treatment resulted in a significant increase in CD8+ T cells but significant reductions of CD4+, γδ and regulatory T cells (Fig 6D). Looking specifically at macrophages, there was a significant enrichment of M1-like macrophages (CD45^+^, CD11b^+^, F4/80^+^, CD86^+^) after combination treatment compared to mono-treatments, while presence of M2-like macrophages (CD45^+^, CD11b^+^, F4/80^+^, CD204^+^) was unaltered (Fig. 6E).

We next investigated the presence of senescent cells in tumor tissue *in vivo* by β-gal staining (Suppl. Fig. 7D). Few senescent cells were visible in vehicle-treated mice, but there was a substantial increase in β-gal+ cells after palbociclib treatment, and some increase after crizotinib treatment, but not to the same extent as after palbociclib treatment. Interestingly, in contrast to treatment of YUMM1.7 cells *in vitro* (Suppl. Fig. 7B), the number of senescent cells decreased after combination treatment compared with palbociclib alone (Suppl. Fig. 7D), suggesting that senescent cells are eliminated after palbociclib + crizotinib-treatment, possibly by infiltrating CD8+ T cells or M1-like macrophages.

To further investigate the influence of CD8+ T cells on the tumor-inhibiting effects of the combination treatment, CD8^+^ T cells were depleted from the mice by injection of α-CD8 antibodies, and the experiments were repeated. Strikingly, CD8+ T cell depletion led to a significant increase in tumor weight compared to control antibodies in palbociclib + crizotinib-treated mice, to a level equivalent to vehicle-treated mice. This suggests that the reduction in tumor weight after combination treatment is at least partly dependent on CD8+ T cells (Fig. 6F).

## Discussion

Targeted therapies and immunotherapy have revolutionized treatment of metastatic melanoma during recent years. Combined ICI therapy results in long-term survival in up to 50% of patients with advanced melanoma (51). In contrast, tumors eventually develop resistance to targeted therapies leading to tumor relapse, emphasizing the urgent need for new treatment strategies (9,52). Many of the genetic aberrations implicated in drug resistance are known to be involved in senescence regulation, and “pro-senescence therapy” or “therapy-induced senescence (TIS)” have therefore been suggested as an alternative therapeutic approach in such tumors (10–12).

Using a selected number of candidate drugs and compounds in a high throughput fluorescence microscopy-based senescence screen we evaluated the TIS strategy in melanoma. Our results showed that the BRAF^V600E^ inhibitor vemurafenib, the MEK1/2 inhibitor trametinib, and the p53-targeting compounds induced senescence in a subset of cell lines. In contrast, the CDK4/6 inhibitor palbociclib, the PI3 kinase inhibitor BKM120 and the MET/ALK/ROS1 inhibitor crizotinib triggered senescence in all or most cell lines, irrespective of *BRAF*/*NRAS* mutation status. CDK4/6 and PI3K inhibitors have previously been shown to induce senescence in different types of tumor cells, including melanoma cells (53–55), but as far as we know, this is the first report that crizotinib can induce senescence in melanoma cells, and there have been very few reports at all relating senescence-induction to this drug in cancer.

Further, our data show that combination of palbociclib, BKM120 or crizotinib with vemurafenib synergistically enhanced senescence and overcame resistance in *BRAF^V600E^*-mutant cell lines, which is in line with previous reports regarding palbociclib and PI3K inhibitors (55–57). Our novel finding that vemurafenib and crizotinib combination treatment synergistically induce senescence in vemurafenib-resistant melanoma cells is interesting considering that activation of the HGF/MET-signaling has been suggested as one escape pathway during development of resistance to BRAF- or MEK-inhibitors in BRAF-mutant melanoma (52,58,59).

The most important novel finding from our work is that the combination of palbociclib and crizotinib synergized to further enhance the senescence response in the melanoma cell lines irrespective of *BRAF*/*NRAS* mutation status. This combination has not been reported in melanoma therapy so far. Interestingly, upregulation of MET signaling has been implicated in resistance to CDK4/6 inhibition in glioblastoma, and dual inhibition of MET/TRK1 and CDK4/6 could overcome this resistance (60). Senescence induced by palbociclib alone and together with crizotinib was associated with expression of SASP factors such as IL-1α, IL-1β, IL-8 and AREG and with increased expression of HLA class I and II (involved in antigen presentation to CD8^+^ and CD4^+^ T-cells, respectively). Palbociclib alone and the combination of palbociclib with crizotinib also increased the expression of PD-L1, which is an immune checkpoint molecule suppressing T-cell and NK cell activation (61,62). This suggests that mono- and combination-treatment with palbociclib and crizotinib induced both positive and negative immune markers with respect to CD8^+^ T cell killing. Moreover, palbociclib and the combination with crizotinib upregulated the expression of the ULBP2/5/6 and MICA/B ligands of the NK cell receptor NKG2D that could both activate NK cells as well as FAS and DR5 death receptors involved in both NK- and T-cell-mediated killing.

Utilizing the syngeneic YUMM1.7 mouse melanoma model, our results showed that palbociclib + crizotinib combination treatment synergistically reduced tumor growth as compared to the mono-treatments. Further, we observed a significant enrichment of CD8+ T cells and M1-like macrophages but reductions of CD4+ T cells including Treg cells as well as of B cells, NK cells and myeloid cells after combination treatment compared with the mono-treatments and with vehicle. The reduction in NK cells was unexpected considering that these cells have been reported to play an important role in clearance of senescent tumor cells in a KRAS-driven mouse model (53). This indicates that CD8+T cells may play a more important role in our model, which is in line with the increased HLA class I expression we observed after treatment. In support of this view, depletion of CD8^+^ T cells resulted in a significant increase in tumor weight, suggesting that the combination of palbociclib and crizotinib indeed enhances CD8^+^ T cell-mediated tumor cell elimination. Interestingly, while the combination treatment increased YUMM1.7 cell senescence in culture, it decreased the number of senescent cells in vivo compared with palbociclib alone, indicating increased clearance of senescent cells *in vivo* in response to combination treatment. While senescence induced by palbociclib and other CDK4/6 inhibitors, alone or in combination with other drugs have been shown to stimulate immune clearance of senescent tumor cells (53,54,63–67), we show here that these effects can be synergistically enhanced by combination treatment with crizotinib in melanoma.

The mechanism behind the synergy between palbociclib and crizotinib with respect to tumor growth inhibition remains to be determined. One possible explanation is that the senescent state becomes more robust and prevents senescence escape pathways more efficiently than single treatments can do. An additional explanation is that the combination treatment reshapes the senescence phenotype in a “senomorphic” way to become more anti-tumorigenic, for instance by modifying the SASP profile or enhancing recognition by the immune cells, thereby affecting their recruitment and activation (10,12,21,25,68–71). It has been reported that treatment with crizotinib in combination with afatinib in melanoma cells downregulates mTOR signaling (38), which is known to affect the composition of SASP (69,70). Furthermore, crizotinib in combination with cisplatin enhanced T cell infiltration and interferon-γ production in a lung cancer model (72), and in another study, MET inhibition blocked recruitment of immunosuppressive neutrophils in a HGF-CDK4^R24C^ -driven mouse melanoma model, thereby improving adoptive T cell transfer or ICI immunotherapy (73). Although MET, rather than ALK or ROS1, is likely the most relevant target of crizotinib in the context of induced senescence in melanoma, this remains to be determined. Clarifying the mechanism(s) by which the combination palbociclib and crizotinib affects the senescence phenotype and immune surveillance in melanoma requires further studies. Another limitation of our study is that it is based on 10 long-term established human melanoma cell lines, and one recently established line. In the future, it would be interesting to investigate the effects of combined palbociclib and crizotinib treatment on a broader set of primary melanoma cultures of different types.

In summary, we found that combination of palbociclib and crizotinib induces senescence, SASP and regulation of immune markers in a broad range of human melanoma cells, including BRAF^V600E^-mutant tumor cells resistant to vemurafenib, NRAS-mutant and non-BRAF/NRAS-mutant melanoma, for which there today are few options for targeted therapy. Importantly, the combination treatment led to a significant reduction of melanoma growth *in vivo*, increase in CD8+ T cells and M1-like macrophages in the tumor microenvironment, and CD8+ T cell-dependent anti-tumor response to treatment. This suggests that palbociclib/crizotinib-based pro-senescence therapy is potentially an alternative treatment strategy for melanoma. The two drugs are already in clinical use for different indications (74,75), which should facilitate further clinical studies of this combination in patients with melanoma. Considering the increased PD-L1 expression in response to palbociclib and crizotinib treatment, further improvement of treatment could be combining these regimens with ICI, or alternatively senolytic drugs to eliminate senescent tumor cells in a “one-two punch” strategy. These will be interesting options to explore in the future with the aim of improving treatment of patients with melanoma.

## Acknowledgements

We thank Dr. Juha Rantala, Misvik Biology, Turku, for advice regarding high-throughput fluorescence microscopy screening. This work was supported by a grant from the Knut and Alice Wallenberg Foundation and by Swedish Cancer Society (Cancerfonden) to LGL.

## Author contribution

SEB, JH, RK, GS, KGW, MW, LGL conceptualized and supervised the study, and acquired funding. FZ, LBo, ID, JM, MSi, MSt, LZ, MA, AA, LB, WB, MDSL, JG, RT, VH, FJ, MW performed experiments, analyzed, visualized and interpreted the data. FZ, LBo, SEB, RK, GS, KGW, MW, LGL wrote the manuscript.

