## Supplemental Information for "Palbociclib CDK4/6- and Crizotinib MET/ALK/ROS1-inhibitors Synergize to Enhance Senescence and Immune Recognition in Melanoma Cells Independently of *BRAF*/*NRAS* Status"

#### Supplemental Materials & Methods

##### Antibodies

Antibodies used in the senescence screen; anti-H3K9me3 (Millipore Cat# 07-523, RRID:AB_310687), anti-p53 (Santa Cruz Biotechnology Cat# sc-126, RRID:AB_628082) and anti-HLA class I (Abcam Cat# ab70328, RRID:AB_1269092).

Antibodies used for western blot analysis:

Primary antibodies included pRbS780 (Cell Signaling Technology Cat# 9307, RRID:AB_330015), Rb (Santa Cruz Biotechnology sc-50, RRID:AB_632339), caspase 3 (Cell Signaling Technology clone 3G2 Cat# 9668, RRID:AB_2069870) and cyclin B (Santa Cruz Biotechnology Cat# sc-595, RRID:AB_2291053, β-actin (Sigma-Aldrich clone AC-15 Cat# A5441, RRID:AB_476744).

Antibodies used for flow cytometry, human cells:

anti-human HLA-ABC APC-Cy7 (BioLegend Cat# 311426, RRID:AB_10709590), CD274 BV786 (BD Biosciences Cat# 563739, RRID:AB_2738397), ULBP2/5/6 PerCP (clone 16590; RRID:AB_3699135), DR5 PE (BioLegend Cat# 307406, RRID:AB_2204926), MICA/MICB APC MICA/MICB APC (Clone 6D4; Biolegend, FAS (BioLegend Cat# 305608, RRID:AB_314546).

Antibodies used for flow cytometry, mouse cells:

anti mouse CD45 BV605 (BioLegend Cat# 103155, RRID:AB_2650656), CD3 APC (BioLegend Cat# 100236, RRID:AB_2561456), CD4 AF488 (BioLegend Cat# 100423, RRID:AB_389302), CD8α PE-Cy7 (BioLegend Cat# 100722, RRID:AB_312761), NK1.1 BV711 (BioLegend Cat# 108702, RRID:AB_313389), γδTCR BV421 (BioLegend Cat# 118120, RRID:AB_2562566), CD19 AF700 (BioLegend Cat# 115528, RRID:AB_493735), CD25 PerCP (BioLegend Cat# 102028, RRID:AB_2295974), FoxP3 PE (BioLegend Cat# 126404, RRID:AB_1089117), CD11b AF700 (BioLegend Cat# 101212, RRID:AB_312795), F4/80 BV421 (BioLegend Cat# 123132, RRID:AB_11203717), CD204 AF488 (Bio-Rad Cat# MCA1322A488, RRID:AB_324818), Ly6G PerCP (BioLegend Cat# 127602, RRID:AB_1089180), CD11c PE-Cy7 (BD Biosciences Cat# 558079, RRID:AB_647251), Ly6C BV510 (BioLegend Cat# 128033, RRID:AB_2562351) and CD86 BV786 (BD Biosciences Cat# 740900, RRID:AB_2740548).

**MUG assay**

For quantitative MUG assay (37), culture media was removed, and cells were washed in PBS. Subsequently, lysis buffer containing 5 mM CHAPS, 40 mM citric acid, 40 mM sodium phosphate and 1x EDTA-free protease inhibitor cocktail was added to wells. Cells were scraped from well surface, collected and vortexed briefly. After spinning samples at 12 000 x g for 5 minutes, supernatants were collected and kept on ice until running the assay. Equal volumes of reaction buffer (pH 6), containing 40 mM citric acid, 40 mM sodium phosphate, 300 mM NaCl, 10 mM beta-mercaptoethanol, 4 mM $MgCl_2$ and 1.7 mM MUG was mixed with 100 µl sample. Bradford assay was used to determine protein concentrations in samples and was subsequently used for normalization of assay data. Samples were incubated at 37°C and at time-points 0 and 90 minutes; a 50 µl sample aliquot was collected. The enzymatic reaction was stopped by diluting the sample aliquots in 0,5 ml 400 mM Sodium carbonate stop solution.

**Primers**

Primers used for the real-time RT-PCR were:

IL-6: FP: CCTGAACCTTCCAAAGATGGC; RP: TTCACCAGGCAAGTCTCCTCA

IL-6R: FP: ATCGGGCTGAACGGTCAAAG; RP: GGCGTCGTGGATGACACAG

IL8: FP: ATGACTTCCAAGCTGGCCGTG; RP: TGTGTTGGCGCAGTGTGGTC

MMP3: FP: AGGGAACTTGAGCGTGAATC; RP: TCACTTGTCTGTTGCACACG

AREG: FP: AGCTGCCTTTATGTCTGCTG; RP: TTTCGTTCCTCAGCTTCTCC

CXCL1: FP: CACCCCAAGAACATCCAAAG; RP: TAACTATGGGGGATGCAGGA

IL1α: FP: CCGTGAGTTTCCCAGAAGAA; RP: ACTGCCCAAGATGAAGACCA

IL1β: FP: CTTCGAGGCACAAGGCACAA; RP: CTGGAAGGAGCACTTCATCTGT

CSF2 (GM-CSF): FP: AAAGGGGATGACAAGCAGAA; RP: ACTACAAGCAGCACTGCCCT

### Supplementary Table 1. Information of drugs and cell lines in the screen.

## A

| | | | | Concentration in µM | | | | | |
|---|---|---|---|---|---|---|---|---|---|
| **Drugs (target)** | **Company** | **Dissolved** | **stock conc.** | **Fold dilution** | **High** | **Mid high** | **Mid** | **Mid low** | **Low** |
| **1.APR-246 (p53)** | Aprea | DMSO | 100 mM | 1.5-fold | 15 | 10 | 6.7 | 4.4 | 2.96 |
| **2.Nutlin 3a (mdm2/p53)** | Selleckchem | DMSO | 10 mM | 3-fold | 10 | 3.333333 | 1.111111 | 0.37037 | 0.123457 |
| **3.RITA (p53)** | NCI | DMSO | 10 mM | 3-fold | 6 | 2 | 0.666667 | 0.222222 | 0.074074 |
| **4.Vemurafenib (BRAF)** | Selleckchem | DMSO | 10 mM | 3-fold | 10 | 3.333333 | 1.111111 | 0.37037 | 0.123457 |
| **5.Trametinib (MEK-inhibitor)** | Selleckchem | DMSO | 10 mM | 3-fold | 0.033 | 0.0111 | 0.0037 | 0.0012 | 0.0004 |
| **6.Crizotinib (ALK, MET)** | Selleckchem | DMSO | 10 mM | 2-fold | 4 | 2 | 1 | 0.5 | 0.25 |
| **7.BKM120 (PI3K)** | Selleckchem | DMSO | 10 mM | 2-fold | 3 | 1.5 | 0.75 | 0.375 | 0.188 |
| **8.Palbociclib (CDK4/6)** | Selleckchem | DMSO | 5 mM | 4-fold | 6 | 1.5 | 0.375 | 0.0938 | 0.0235 |
| **9. Seliciclib (CDK2)** | Selleckchem | DMSO | 10 mM | 2-fold | 30 | 15 | 7.5 | 3.75 | 1.895 |
| **10. Temozolomide (DNA alkylator)** | Selleckchem | DMSO | 118 mM | 2-fold | 600 | 300 | 150 | 75 | 37.5 |

## B

| Cell line | Cell number at seeding | Culture medium |
|---|---|---|
| **A375** | 900 | DMEM |
| **KADA** | 600 | DMEM |
| **SKMEL28** | 1200 | MEM |
| **SKMEL2** | 1500 | MEM |
| **ESTDAB37** | 800 | RPMI |
| **ESTDAB102** | 1500 | RPMI |
| **ESTDAB105** | 1250 | RPMI |
| **ESTDAB49** | 800 | RPMI |
| **ESTDAB138** | 1250 | RPMI |
| **ESTDAB140** | 1250 | RPMI |

(A) Information of drugs regarding to company, solvent, stock concentration, fold for serial dilutions and drug concentrations as high (H), mid high (MH), mid (M), mid low (ML) and low (L) dose used both in the screen and in the validation. (B) Cell density for seeding in 384-well plate and optimal media for individual cell lines.

#### Supplementary Table 2. Highest starting concentrations during titration of drugs in the combination senescence screen.

##### A VEM combo

| Conc. (µM) | VEM | TMZ | BKM | CRIZ | PAL | TRA |
|---|---|---|---|---|---|---|
| A375 | 10 | 600 | 3 | 10 | 5 | |
| A375 | 0.156 | | | | | 0.625 |
| A375-VR4 | 5 | 600 | 3 | 10 | 5 | 0.625 |
| ESTDAB37 | 5 | 600 | 3 | 10 | 3 | 0.625 |

##### B PAL combo

| Conc. (µM) | PAL | BKM | CRIZ |
|---|---|---|---|
| A375/VR4/ESTDAB37 | 5 | 1.5 | 2.5 |

##### C TRA combo

| Conc. (µM) | TRA | TMZ | BKM | CRIZ | PAL | APR | RITA | F4 |
|---|---|---|---|---|---|---|---|---|
| KADA | 10 | 600 | 10 | 10 | 5 | 40 | 50 | 100 |

(A) in the vemurafenib combination; (B) in palbociclib combination; (C) in trametinib combination.

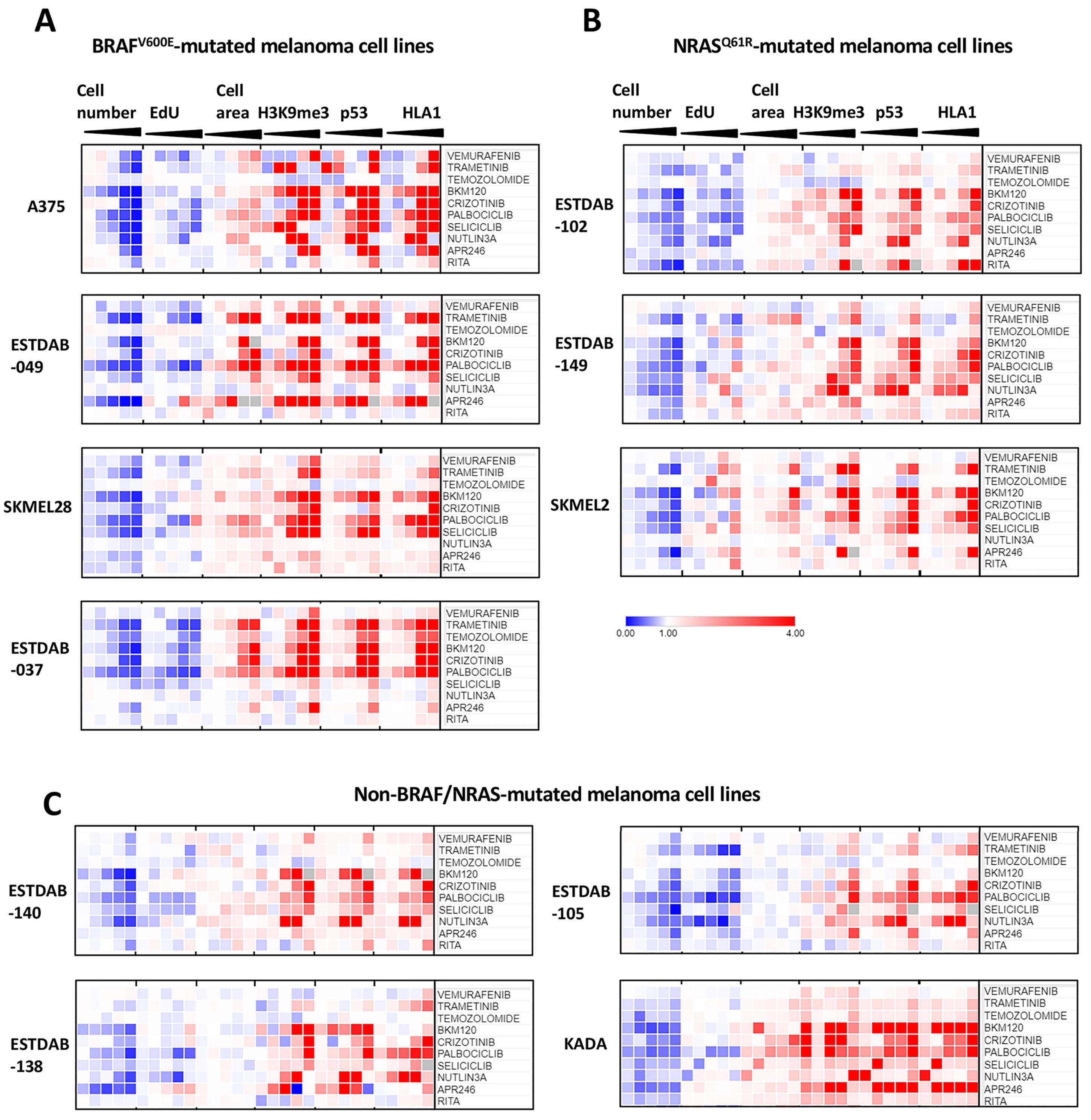


**Supplemental Figure 1**

**Supplementary Figure S1**. Screen for drug-induced senescence in human melanoma cell lines. (A-C) Heatmaps showing data from measurements of six biomarkers in response to 11 selected drugs after 72 hours of treatment. Drug concentrations are specified in Supplementary Table 1A, and increasing concentrations are illustrated as wedges. (A) Four melanoma cell lines with $BRAF^{V600E}$ mutation; (B) Three melanoma cell lines with $NRAS^{Q61R}$ mutation; (C) Four melanoma cell lines with non-BRAF/NRAS mutations. Intensity per field was normalized to DMSO and median values (%) were presented here with indicated color code.

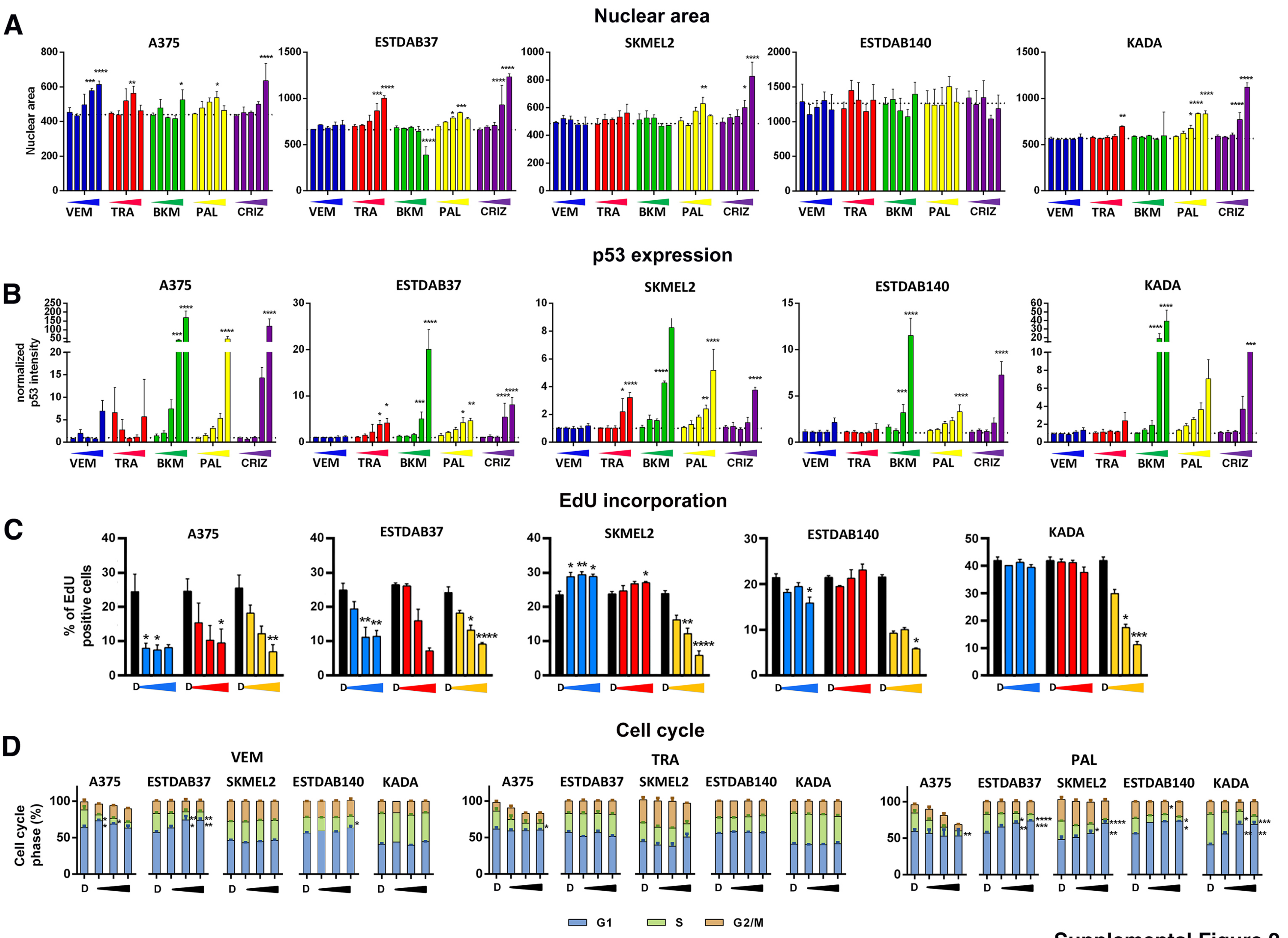


Supplemental Figure 2

**Supplementary Figure S2.** Validation of senescence induction in response to selected drugs in selected melanoma cell lines. Single-drug treatment of A375, ESTDAB37, SKMEL2, ESTDAB140 and KADA with vemurafenib (VEM), trametinib (TRA), BKM120 (BKM), palbociclib (PAL) or crizotinib (CRIZ) for 3 days. (A and B) Bar charts illustrating nuclear size (A) and intensity of p53 staining (B). Drug concentrations are specified in Supplementary Table 1A. Data was normalized to DMSO treatment and mean±SEM values are presented. The value for DMSO for each marker is indicated as a dotted line in the figure. (A-C) Blue: VEM; red: TRA; green: BKM; yellow: PAL; purple, CRIZ. Increasing concentration are illustrated as wedges. (C) Percentage of cells incorporating EdU as determined by flow cytometry. Data are presented as Mean±SEM. (D) Cell cycle analysis as determined by flow cytometry. Blue: G1 phase; green: S phase; orange: G2/M phase. (C and D) Drugs concentrations are mid-high, mid and mid-low, respectively (see Supplementary Table 1A), as illustrated as wedges. D=DMSO. * $p< 0.05$; ** $p< 0.01$, *** $p<0.001$, **** $p<0.0001$.

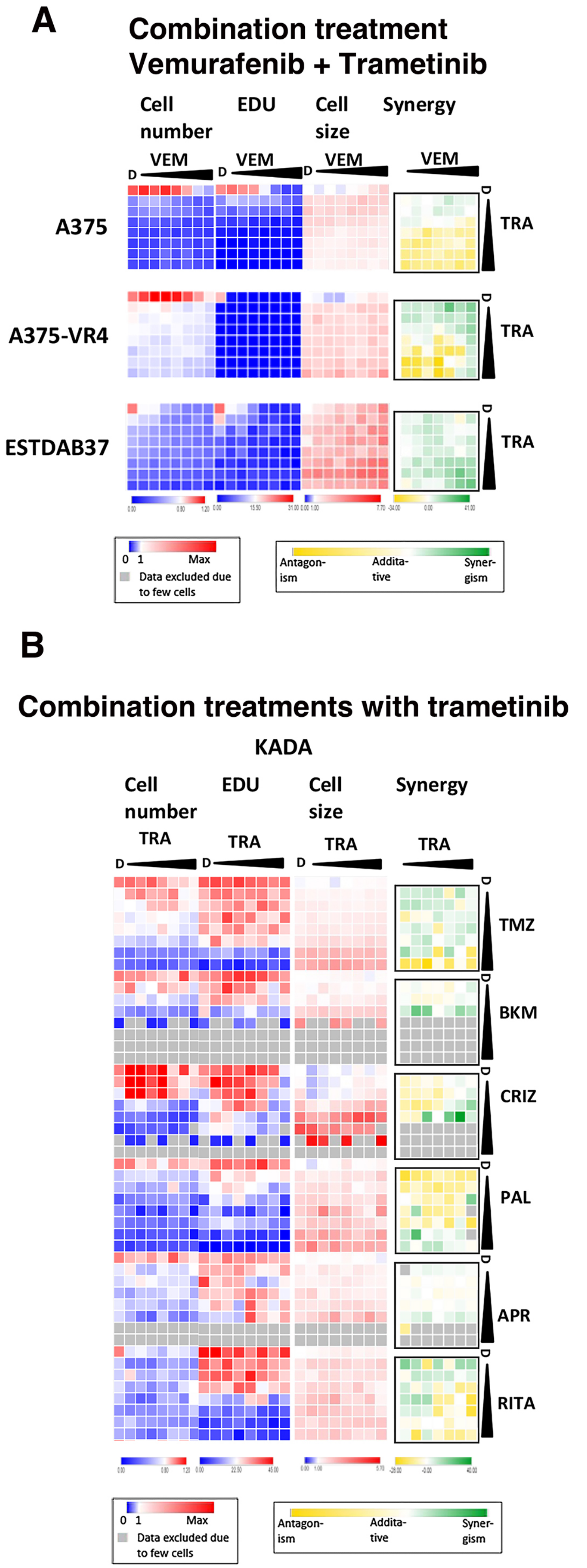


Supplemental Figure 3

**Supplementary Figure S3**. (A) Drug combination senescence screen based on vemurafenib and trametinib using selected melanoma cell lines. Heatmaps of cell number, % EdU positive cells, cell area and synergy calculation of cell area on A375, A375-VR4 (acquired VEM-resistant) and ESTDAB37 (intrinsically VEM-resistant) cells upon the combination of VEM and TRA treatment. Indicated cell lines were treated with the drugs at 2-fold serial dilutions for 72 hours. The highest doses of the drugs are shown in Suppl. Table 2, and increasing concentration are illustrated as black wedges. D=DMSO. Cell number and cell size were normalized to DMSO-treated cells. (B) Drug combination senescence screen based on the MEK1/2 inhibitor trametinib (TRA). Heatmaps of cell number, % EdU positive cells, cell area and synergy calculation of cell area on KADA cells upon the combination of TRA with temozolomide (TMZ), BKM120 (BKM), crizotinib (CRIZ), palbociclib (PAL), APR-246 (APR) or RITA. The cells were treated with the drugs at 2-fold serial dilutions for 72 hours. The highest doses of the drugs are shown in Suppl. Table 2, and increasing concentration are illustrated as black wedges. D=DMSO. Cell number and cell size were normalized to DMSO-treated cells. Missing data due to too low cell number as a result of cell death is indicated as grey. Green = synergism, yellow = antagonism.

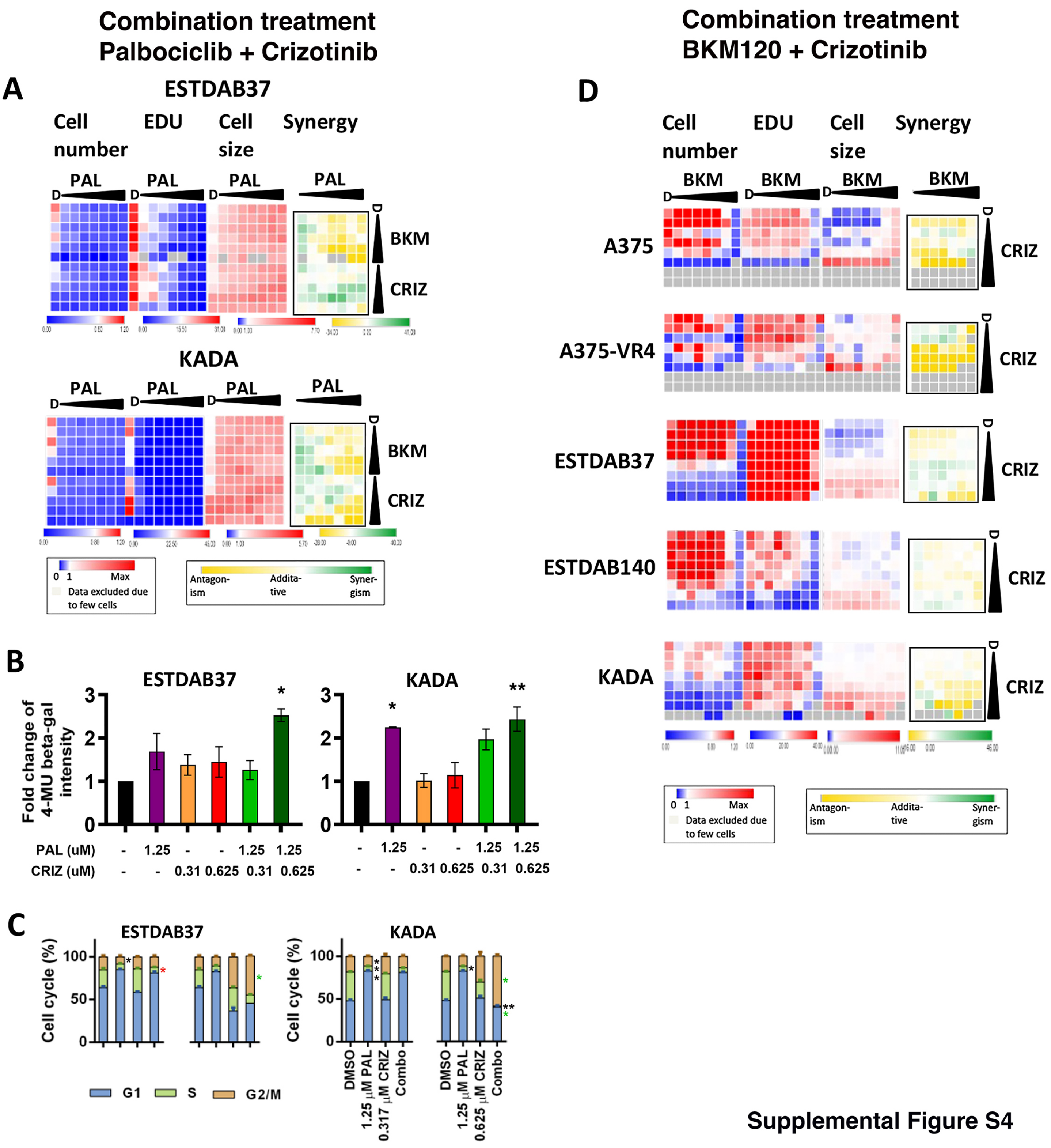


Supplemental Figure S4

**Supplementary Figure S4.** Drug combination senescence screen using selected melanoma cell lines. (A) Combination treatment using PAL together with BKM or CRIZ at 2-fold serial dilutions for 72 hours. The highest doses of the drugs are listed in Suppl. Table 2B. The heatmaps show data representing cell number, percentage of EdU positive cells, cell size and combination synergy index based on cell size in ESTDAB37 and KADA cells. Missing data due to too low cell number as a result of cell death is indicated as grey. Green = synergism, yellow = antagonism. (B) Fluorescent β-galactosidase assay (MUG) after mono or combination treatment with PAL and CRIZ as indicated. (C) Cell cycle analysis after mono or combination treatment with PAL and CRIZ in indicated cell lines for 72 hours. * $p < 0.05$; ** $p < 0.01$. Black stars represent significance DMSO control vs. treatments, red and green stars represent significance CRIZ vs. combination and PAL vs. combination treatments, respectively. (D) Drug combination senescence screen based on BKM120 and crizotinib using selected melanoma cell lines. Heatmaps of cell number, % EdU positive cells, cell area and synergy calculation of cell area. Indicated cell lines were treated with BKM together CRIZ at 2-fold serial dilutions for 72 hours. The highest doses of the drugs are shown in Suppl. Table 2, and increasing concentration are illustrated as black wedges. D=DMSO. Cell number and cell size were normalized to DMSO-treated cells. Missing data due to too low cell number as a result of cell death is indicated as grey. Green = synergism, yellow = antagonism.

### SASP expression

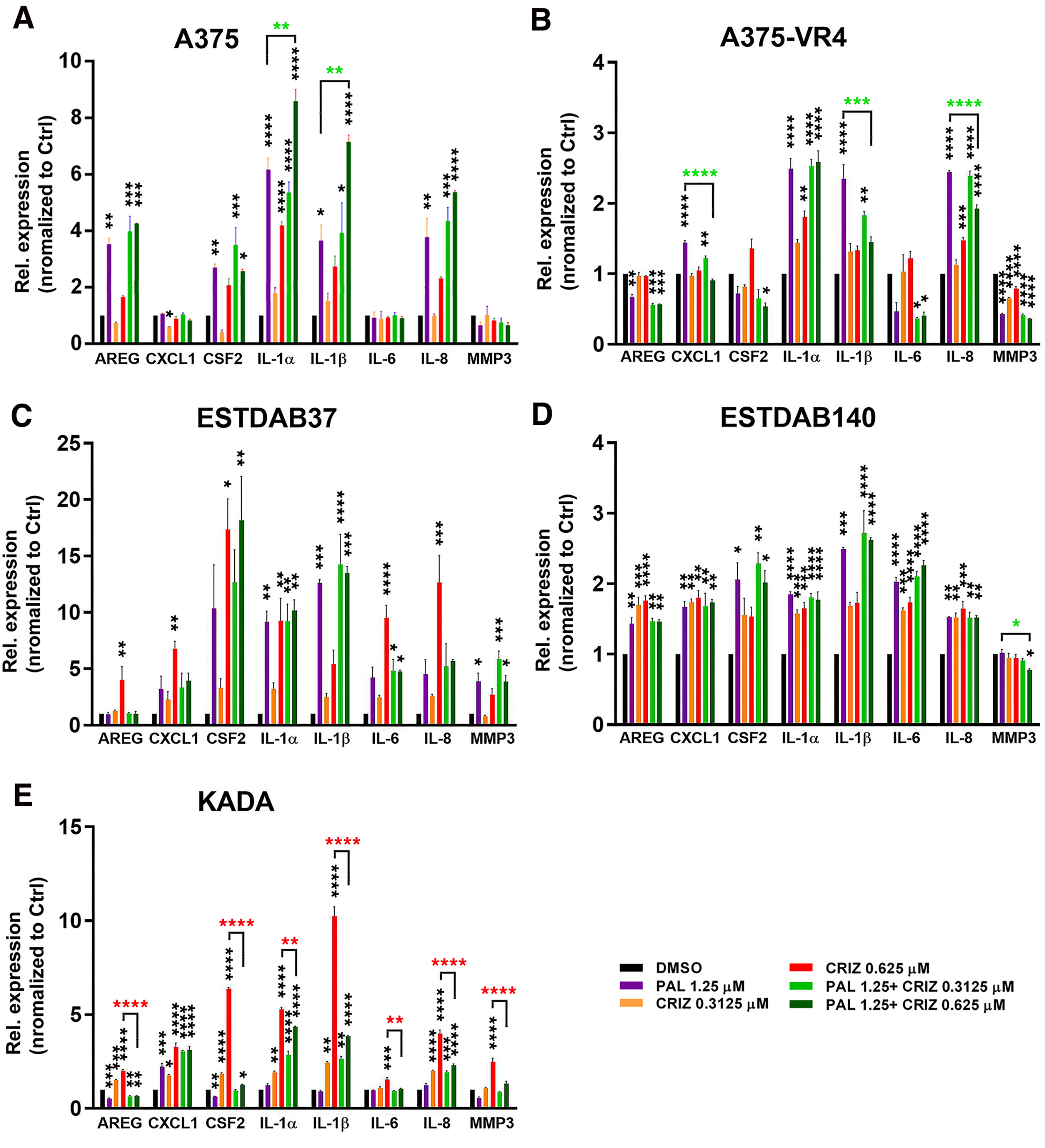


Supplemental Figure 5

**Supplementary Figure S5.** SASP-related gene expression in response to combination treatment using palbociclib (PAL) and crizotinib (CRIZ). Relative fold change of SASP-related gene expression in selected melanoma cells lines upon indicated treatments for 72 hours. (A) A375 (results for all 8 genes, including those already presented in Fig. 4D) (B) A375-VR4 (C) ESTDAB37 (D) ESTDAB140 and (E) KADA cells. Biological triplicates were presented as mean±SEM and one-way ANOVA was applied to compare drug treatment versus mono-therapy. * $p < 0.05$; ** $p < 0.01$, *** $p < 0.001$, **** $p < 0.0001$. Black stars represent significance DMSO control vs. treatments and are placed above columns in the graphs, red and green stars represent significance CRIZ vs. combination and PAL vs. combination treatments, respectively, and are placed above brackets between columns.

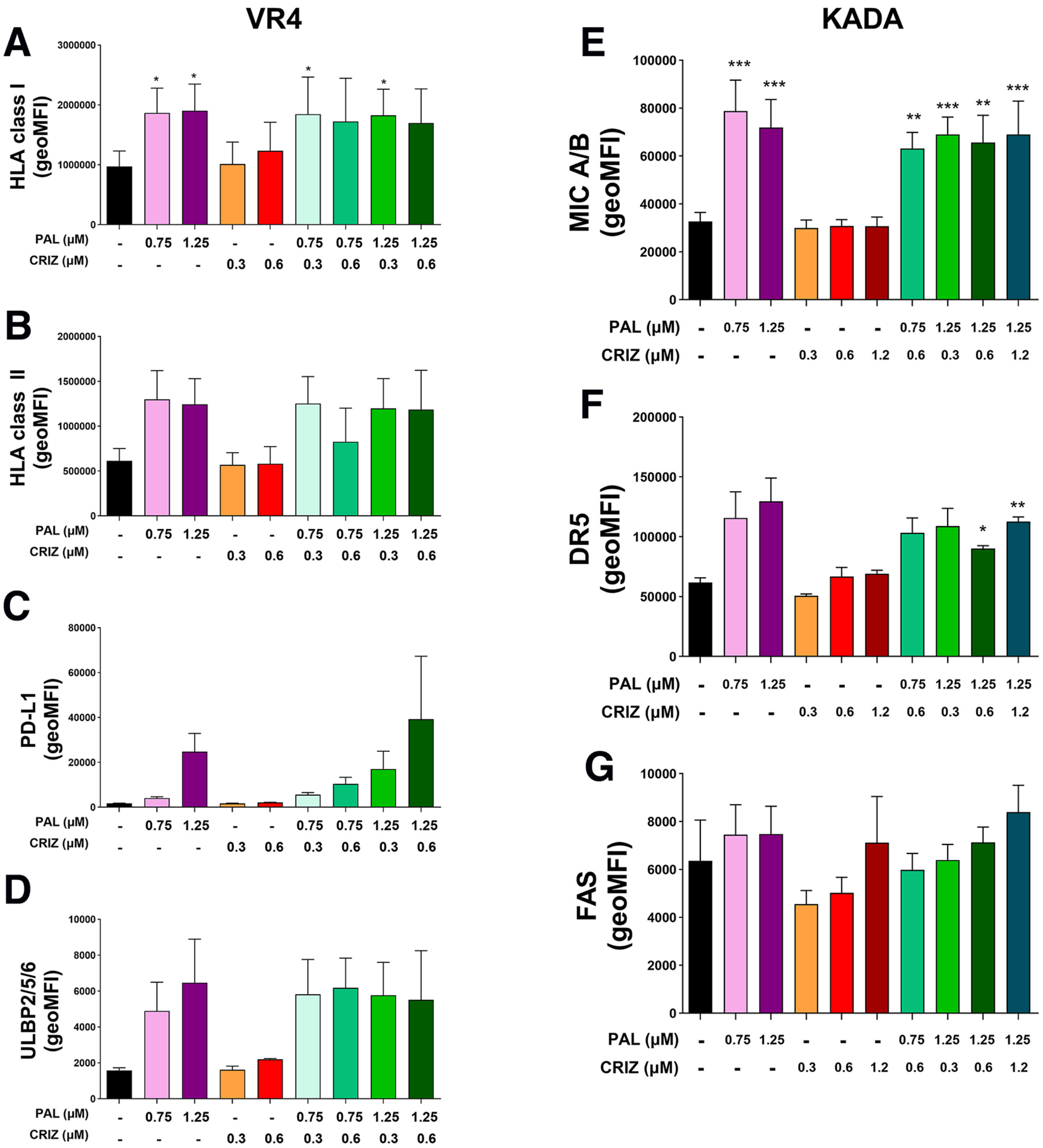


Supplemental Figure 6

**Supplementary Figure S6.** Expression of immune markers regulating recognition by T cells and NK cells upon combination treatment with palbociclib and crizotinib. (A-D) A375-VR4 cells were treated with indicated concentrations of PAL and/or CRIZ for 72 hrs followed by analysis of (A) HLA class I, (B) HLA class II, (C) PD-L1, (D) ULBP2/5/6. (E-G) KADA cells were treated with indicated concentrations of PAL and/or CRIZ for 72 hrs followed by analysis of (E) MICA/B, (F) DR5 and (G) FAS by flow cytometry using specific antibodies. * $p < 0.05$; ** $p < 0.01$, *** $p < 0.001$, **** $p < 0.0001$. Stars placed above columns in the graphs represent significance DMSO control vs. treatments, and stars placed above brackets between columns represent significance mono-treatment vs. combination treatments.

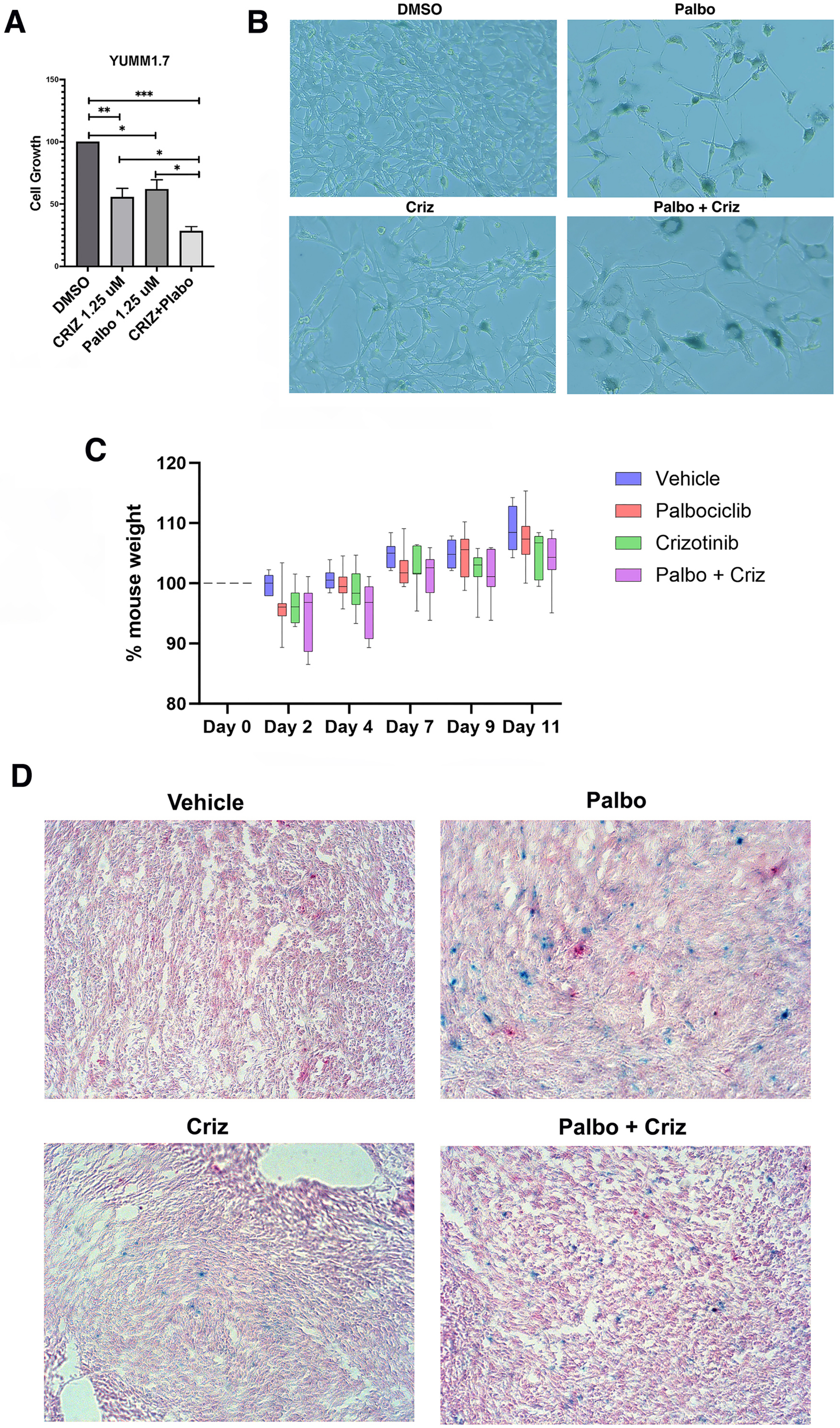


Supplemental Figure 7

**Supplementary Figure S7.** Cell growth and senescence of YUMM1.7 cells in response to palbociclib (Palbo) and crizotinib (Criz) treatment alone or in combination in culture. (A) YUMM1.7 cell growth in response to 72 hours of treatment with Palbo, Criz, or their combination as indicated as determined by resazurin assay. (B) β-gal staining of YUMM1.7 cells after palbociclib (0.625 μM) and crizotinib (5 μM) treatment alone or in combination for 7 days. (C) Effects of mono and combination treatment using Palbo and Criz on C57BL6 mouse weights measurement during treatment period (11 days), corresponding to the experiment shown in Fig. 6A-E. (D) β-gal staining of YUMM1.7 tumor tissue at endpoint after 11 days mono or combination treatments with Palbo and Cris. * $p < 0.05$; ** $p < 0.01$, *** $p < 0.001$.
